# A neural code for egocentric spatial maps in the human medial temporal lobe

**DOI:** 10.1101/2020.03.03.973131

**Authors:** Lukas Kunz, Armin Brandt, Peter C. Reinacher, Bernhard P. Staresina, Eric T. Reifenstein, Christoph T. Weidemann, Nora A. Herweg, Melina Tsitsiklis, Richard Kempter, Michael J. Kahana, Andreas Schulze-Bonhage, Joshua Jacobs

## Abstract

Spatial navigation relies on neural systems that encode information about places, distances, and directions in relation to the external world or relative to the navigating organism. Since the proposal of cognitive maps, the neuroscience of navigation has focused on allocentric (world-referenced) neural representations including place, grid, and head-direction cells. Here, using single-neuron recordings during virtual navigation, we identify “anchor cells” in the human brain as a neural code for egocentric (self-centered) spatial maps: Anchor cells represent egocentric directions towards “anchor points” located in the environmental center or periphery. Anchor cells were abundant in parahippocampal cortex, supported full vectorial representations of egocentric space, and were integrated into a neural memory network. Neurons encoding allocentric direction complemented anchor-cell activity, potentially assisting anchor cells in transforming percepts into allocentric representations. Anchor cells may facilitate egocentric navigation strategies, may support route planning from egocentric viewpoints, and may underlie the first-person perspective in episodic memories.

## Introduction

Spatial navigation and spatial memory are vital for the survival of humans and animals (Ekstrom et al., 2018). Our daily life critically relies on the ability to remember familiar places, to retrieve and to navigate to goal destinations, and to explore and encode new environments. When aging or neurological diseases affect the neurobiological mechanisms underlying these processes, symptoms of spatial disorientation arise and make it difficult for the patients to perform activities of daily living (Coughlan et al., 2018; Ekstrom et al., 2018).

Humans and animals orient themselves and navigate by representing information about places, distances, and directions in different reference frames: in allocentric reference frames that are bound to the external world or in egocentric reference frames that are centered on the navigating subject (Figure S1) (Klatzky, 1998). Behavioral studies have disentangled the relative contributions of allocentric and egocentric spatial representations, which complement each other to support efficient spatial behavior in everyday life (Burgess, 2006; Ekstrom and Isham, 2017; Ekstrom et al., 2014; Meilinger and Vosgerau, 2010; Waller and Hodgson, 2006; Wang and Spelke, 2002; Zhang et al., 2014).

Traditionally, the neuroscience of spatial navigation has focused on the neural codes that underlie allocentric spatial representations (Epstein et al., 2017; Kunz et al., 2019a; Moser et al., 2017): a place or grid cell may indicate if an animal or human is in the “northeast” corner of an environment (Ekstrom et al., 2003; Hafting et al., 2005; Ismakov et al., 2017; Jacobs et al., 2013; O’Keefe and Dostrovsky, 1971), a head-direction cell may activate whenever navigating “south” (Taube et al., 1990), and a boundary vector/border cell may respond to a spatial boundary located to the “west” (Lever et al., 2009; Solstad et al., 2008). Together, these single-neuron codes provide the navigating organism with a mental map of the spatial environment that encodes spatial information in allocentric coordinates (Tolman, 1948; McNaughton et al., 2006; Bellmund et al., 2018; Behrens et al., 2018).

The neural basis of egocentric spatial representations in humans has, however, remained unknown. Hence, in the present study, we addressed this critical gap of knowledge and hypothesized that neurons in the human medial temporal lobe keep track of the instantaneous egocentric relationship between the navigating subject and proximal areas of the environment. Specifically, we tested for the existence of human neurons (“anchor cells”) whose activity encodes the subject’s egocentric direction towards local reference points (“anchor points”). Such a coding scheme would be instrumental for egocentric navigation, because it represents the proximal spatial layout relative to the subject’s viewpoint, which provides self-centered orientation and allows the planning of routes from a first-person perspective.

By demonstrating that humans have anchor cells with anchor points in various locations of the environment, that these cells support full vectorial representations of egocentric space, and that their firing characteristics change as a function of spatial memory performance, we provide the first evidence for a behaviorally relevant single-neuron substrate of egocentric spatial maps in the human medial temporal lobe.

## Results

### Single-neuron recordings in epilepsy patients performing a virtual spatial navigation task

To detect and characterize human anchor cells, we recorded single-neuron activity from the medial temporal lobe of 14 neurosurgical epilepsy patients (Table S1), while they performed an object–location memory task in a virtual environment (Figures 1A and 1B). In this task (Kunz et al., 2015, 2019b), patients learned and repeatedly retrieved the locations of eight different objects. While patients freely navigated the virtual environment throughout the task, their virtual heading directions and locations were recorded to associate these variables with single-neuron activity. Patients contributed a total of 18 sessions (duration between 22 and 74 min) and completed between 34 and 167 trials per session. Patients performed the task well, as spatial memory performance was above chance on 83% of the trials (Figure 1C) and performance increased over the course of the session (Wilcoxon signed-rank test, *z* = 2.896, *P* = 0.004; Figure 1D). Across all sessions, we recorded a total of *N* = 729 neurons from amygdala (*n* = 242), entorhinal cortex (*n* = 114), fusiform gyrus (*n* = 25), hippocampus (*n* = 146), parahippocampal cortex (*n* = 65), temporal pole (*n* = 128) and visual cortex (*n* = 9; Figures S2 and S3).

**Figure 1.**
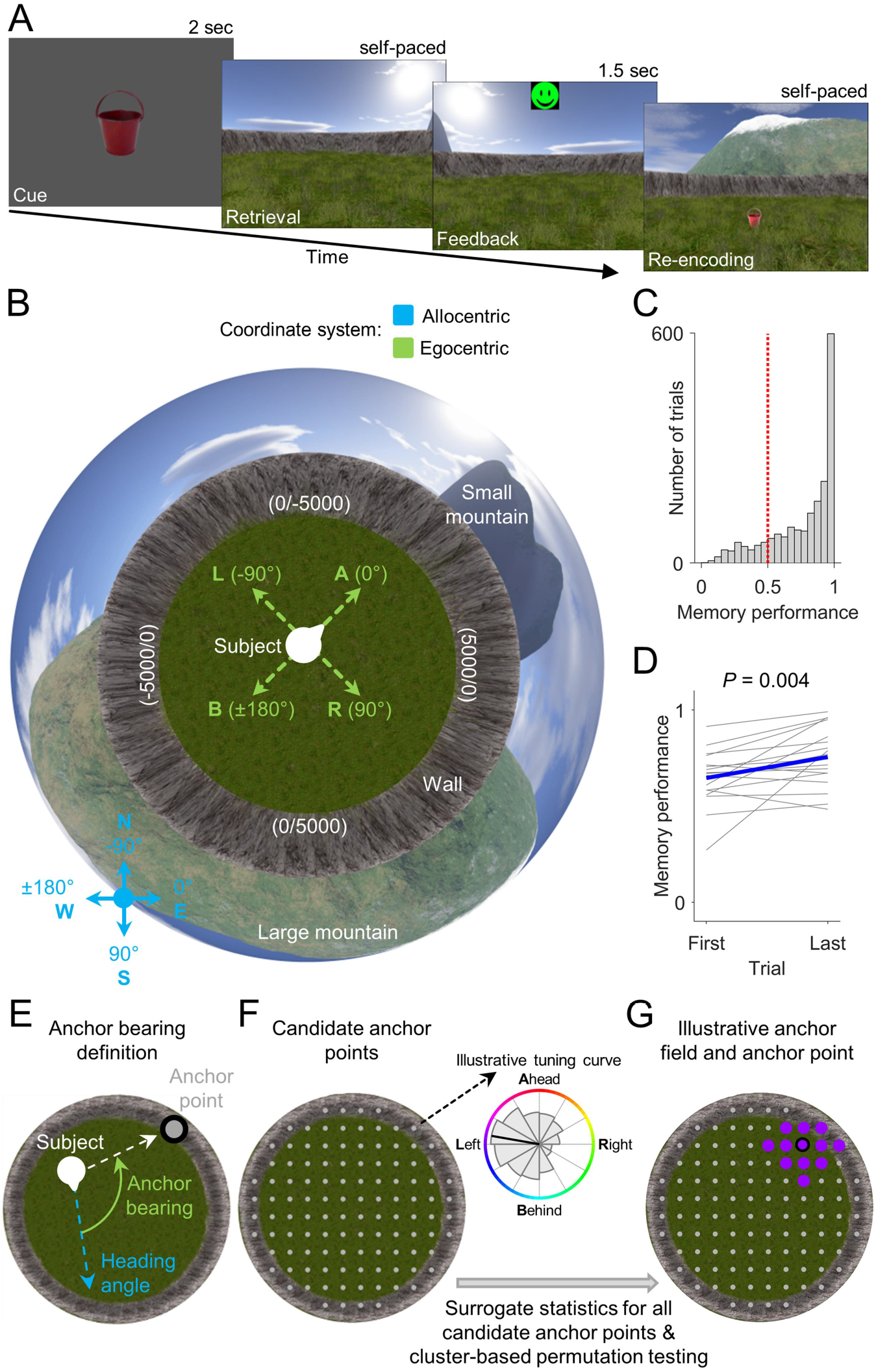
Task, behavioral performance, and illustration of anchor-cell identification. (**A**) Patients performed an object–location memory task in a virtual environment. In each trial, a given object (“cue”) had to be placed at its correct location (“retrieval”). Patients received feedback depending on response accuracy (“feedback”) and re-encoded the object location afterwards (“re-encoding”). (**B**) Bird’s eye view of the environment. Blue compass, allocentric direction; green compass, egocentric direction (given the patient is facing “northeast”). A (B; L; R), ahead (behind; to the left; to the right) of the subject. (**C**) Histogram of spatial memory performance values across all trials from all patients. Red dotted line, chance level. (**D**) Change in spatial memory performance between the first and the last trial. Blue line, mean across subjects. (**E**) Illustration of anchor bearing, which is the angular difference between the allocentric heading angle and the angle of the vector from the subject’s location to the anchor point. (**F**) Left, candidate anchor points. Right, illustrative tuning curve for one candidate point depicting firing rate as a function of bearing towards this point. Significance of each candidate anchor point is tested via surrogate statistics. (**G**) Cluster-based permutation testing identifies the largest cluster of significant candidate anchor points (“anchor field”). The “anchor point” is the center of mass of the anchor field.

### Anchor cells encode egocentric directions towards local reference points

We identified anchor cells by analyzing each neuron’s firing rate as a function of the patient’s egocentric direction (“bearing”) towards local reference points in the virtual environment (Figure 1E; Figures S4 and S5; STAR Methods; Supplemental Text S1). Briefly, for each cell we iterated through 112 candidate anchor points (Figure 1F), each time assessing the degree to which the cell’s firing rate varied as a function of the subject’s egocentric bearing towards this candidate anchor point. The center of mass of the largest cluster of significant candidate anchor points (“anchor field”) defined the anchor point (Figure 1G). This analysis procedure thus resulted in the identification of individual neurons that behaved as anchor cells by tracking the subject’s bearing towards their anchor points within the virtual environment.

We observed 95 anchor cells, 13.0% of all neurons, which is significantly more than expected by chance (binomial test versus 5% chance, *P* < 0.001). On average, there were 5.3 ± 1.3 (mean ± SEM) anchor cells per session. At least one anchor cell was found in 16 of 18 sessions and in twelve of 14 patients. Control analyses showed a significant proportion of anchor cells after excluding neurons from the anchor-cell analysis that were recorded on microelectrodes located in brain regions potentially involved in the generation of epileptic seizures as defined by clinical criteria (80 anchor cells of 570 cells; 14.0%; binomial test, *P* < 0.001).

Anchor cells exhibited anchor points in various locations of the environment and showed a range of preferred egocentric bearings towards these anchor points (Figure 2). For example, the anchor cell in Figure 2A had its anchor point in the “northeast” part of the environment (Figure 2A, left), and the cell’s firing rate increased when this anchor point was ~45° to the right of the subject’s current heading (Figure 2A, middle). We illustrate this cell’s tuning towards its anchor point by plotting the cell’s preferred allocentric direction as a function of location (Figure 2A, right). For anchor cells, this vector-field map often exhibited a systematic change in the cell’s preferred allocentric direction across the environment. Such a pattern is fundamentally different from the homogeneous vector-field maps of neurons encoding allocentric direction (see below).

**Figure 2.**
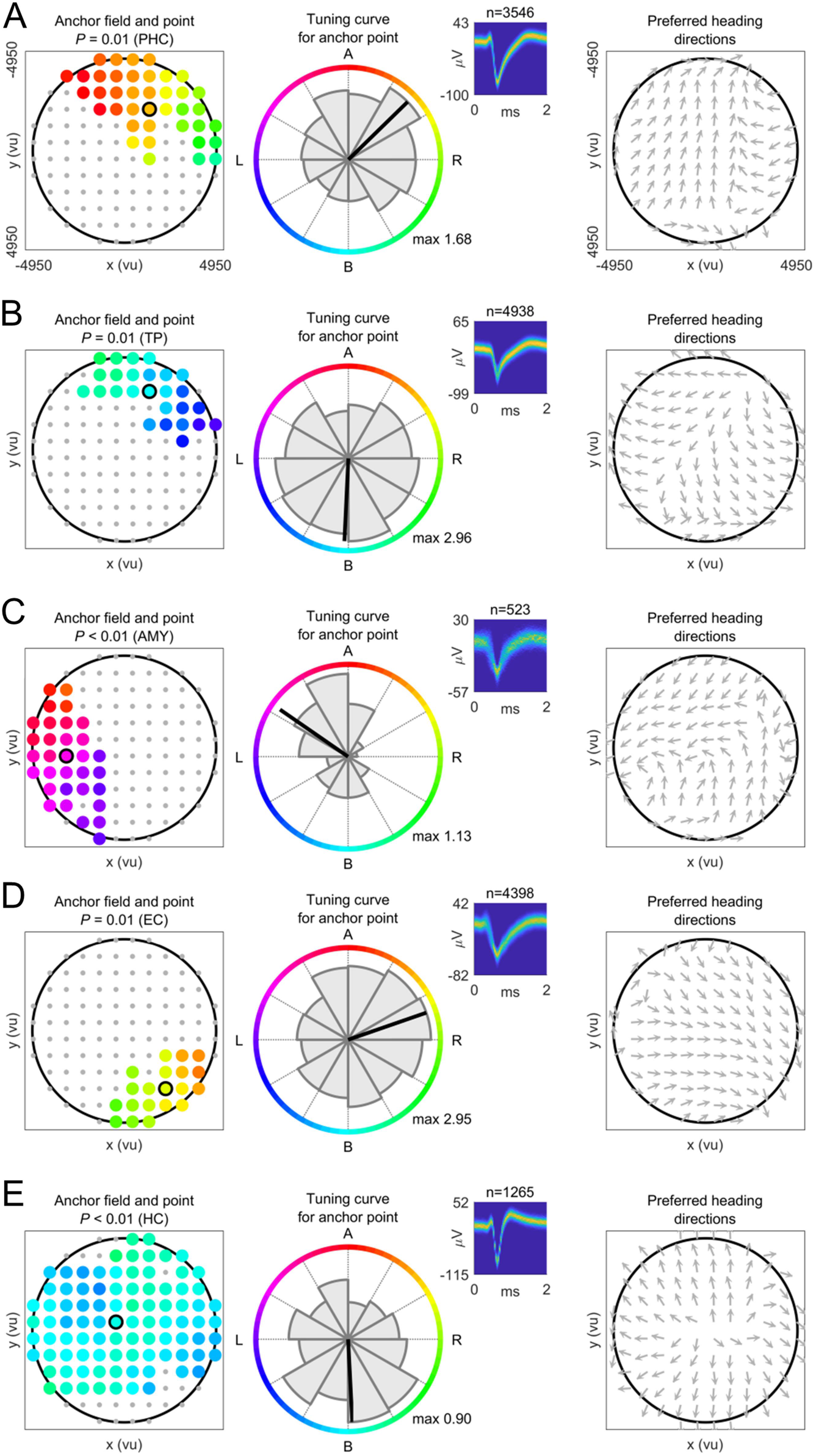
Anchor cells encode egocentric directions towards local reference points (anchor points). (**A to E**) Examples of anchor cells. Left column, anchor-cell plot showing the anchor field (colored dots) and the anchor point (colored dot with black circle). Coloring depicts the preferred egocentric bearing towards each location of the anchor field. The color code corresponds to the colored circle in the middle column. For example, a green-colored dot indicates that the cell activates when this location is to the right of the subject. Gray dots, non-significant candidate anchor points. Large black circle, environmental boundary. *P*-value, significance from cluster-based permutation testing. Middle column, tuning curve showing how the cell’s firing rate varies as a function of egocentric bearing towards the anchor point; max, maximum firing rate (Hz). Colored circle indicates bearing. Inset depicts spike-density plot (number above inset indicates spike number entering the analysis). Right column, vector-field map showing the cell’s preferred allocentric heading direction across the environment (gray arrows). Black circle, environmental boundary. A (B; L; R), anchor point ahead (behind; to the left; to the right) of the subject. ms, milliseconds; vu, virtual units. AMY, amygdala; EC, entorhinal cortex; HC, hippocampus; PHC, parahippocampal cortex; TP, temporal pole.

### Basic properties of anchor cells

Anchor cells were most prevalent in parahippocampal cortex (30.8%; Figure 3A), which is the human homologue of the rodent postrhinal cortex (Aminoff et al., 2013). Some anchor cells showed additional firing-rate modulations related to the patients’ allocentric direction or location (Figures 3B and S6), but a significant number of “pure” anchor cells remained after excluding anchor cells that were also direction and/or place cells (*n* = 60; 8.2%; binomial test, *P* < 0.001).

**Figure 3.**
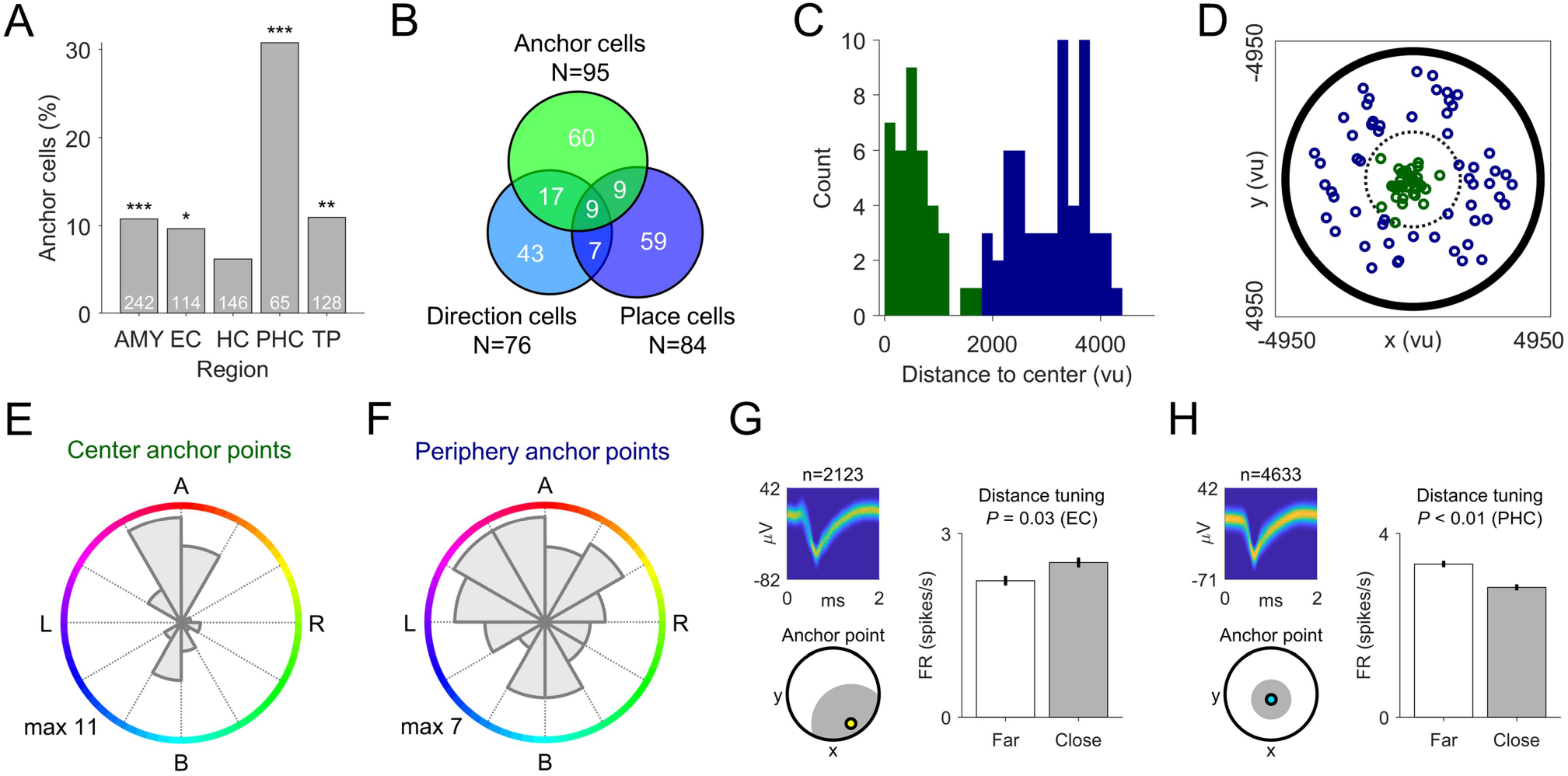
Anchor cells have anchor points in various locations, show a range of egocentric bearings, and exhibit distance tuning. (**A**) Distribution of anchor cells across brain regions. (**B**) Overlap between anchor cells, direction cells, and place cells. (**C**) Distribution of anchor-point distances to the environmental center. Green, center anchor points; blue, periphery anchor points. (**D**) Spatial distribution of anchor-point locations. Black dotted line separates center anchor points (green) from periphery anchor points (blue). (**E**) Distribution of preferred bearings towards anchor points in the environmental center. (**F**) Distribution of preferred bearings towards anchor points in the environmental periphery. (**G and H**) Example anchor cells showing activity correlated with anchor-point distance. Bottom left subpanel indicates the anchor point, with close (gray) and far (white) locations. A (B; L; R), anchor point ahead (behind; to the left; to the right) of the subject. AMY, amygdala; EC, entorhinal cortex; HC, hippocampus; PHC, parahippocampal cortex; TP, temporal pole. max, maximum number; ms, milliseconds; vu, virtual units. Error bars indicate SEM. \**P* < 0.05; \*\**P* < 0.01; \*\*\**P* < 0.001.

Across cells, anchor points were positioned in many different locations of the environment, including both the center and the periphery (Hartigan’s dip test, *P* < 0.001; Figures 3C and 3D). The anchor cells with anchor points in the center of the environment are potentially related to the previously identified center-bearing cells (LaChance et al., 2019) and path cells (Jacobs et al., 2010). The preferred bearings of anchor cells with anchor points in the center of the environment showed a bimodal distribution with an overrepresentation of “ahead” and “behind” bearings (Rayleigh test for two-fold symmetry, *z* = 13.619, *P* < 0.001; Figure 3E), whereas anchor cells with anchor points in the periphery showed a roughly uniform distribution of preferred anchor bearings (Figure 3F).

### Anchor cells support full vectorial representations of egocentric space

To reveal whether anchor cells support a full vectorial representation of egocentric space, we next tested for anchor cells that represented the distance towards the anchor point, in addition to bearing. Hence, for time periods when the subject’s current anchor bearing was in alignment (±90°) with the cell’s preferred anchor bearing, we examined whether the cell’s activity represented the subject’s anchor-point distance. Indeed, 23 of 95 anchor cells showed positive or negative firing-rate changes according to the patient’s distance to the anchor point (13 and 10, respectively; binomial tests, both *P* < 0.028; for examples, see Figures 3G and 3H), comparable to the prevalence of distance tuning found in other spatial cell types (LaChance et al., 2019; Wang et al., 2018). This kind of full vectorial representation of space in egocentric coordinates could be useful for navigation by allowing the navigating organism not only to estimate the direction, but also to compute the location of the anchor point relative to itself.

### Allocentric direction cells complement anchor-cell functioning

Theoretical accounts of spatial navigation and memory suggest that neural transformation circuits combine egocentric spatial representations with allocentric direction information in order to create allocentric spatial representations such as place cells (Bicanski and Burgess, 2018; Wang et al., 2020). Thus, after demonstrating anchor cells that represent spatial information in egocentric coordinates, we tested for single-neuron codes of allocentric direction. We identified 76 “direction cells”, which showed increased firing when patients were heading towards specific global directions (binomial test, *P* < 0.001). For example, direction cells activated when the subject was moving “west” (Figures 4A and 4B). As expected, direction cells exhibited more consistent directional tuning across the environment than anchor cells (i.e., more homogeneous vector-field maps; Wilcoxon rank-sum test, z = 5.392, *P* < 0.001; Figures 4C and 4D), illustrating the difference in their coding schemes (Supplemental Text S2).

**Figure 4.**
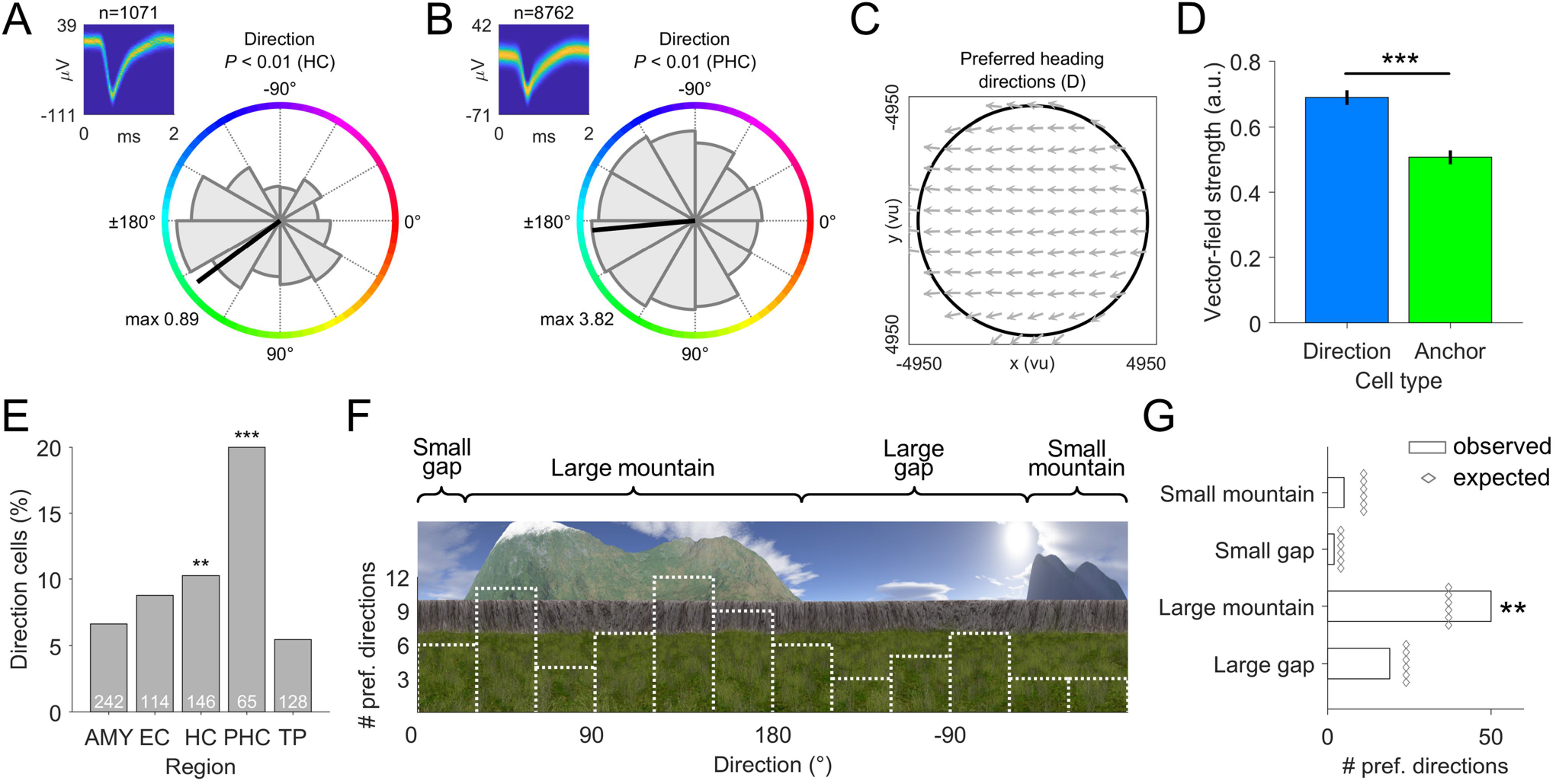
Direction cells complement anchor-cell functioning by representing allocentric direction. (**A and B**) Examples of direction cells encoding allocentric direction. Gray shaded area, tuning curve; black line, preferred direction; colored circle, allocentric direction; max, maximum firing rate (Hz). Upper left subpanels show spike-density plots (number above subpanel indicates spike count). (**C**) Example vector-field map of the direction cell shown in (**B**), illustrating that allocentric direction tuning is similar across the environment. (**D**) Comparison of vector-field strengths between direction cells and anchor cells. (**E**) Distribution of direction cells across brain regions. (**F**) Distribution of preferred directions of allocentric direction cells (white dotted line), shown relative to the background scenery in the virtual environment. (**G**) Bar graph showing the number of preferred directions tuned towards different distal landmarks of the environment’s background. White bars, observed number of preferred directions oriented towards a specific distal landmark; gray diamonds, number of preferred directions expected by chance. AMY, amygdala; EC, entorhinal cortex; HC, hippocampus; PHC, parahippocampal cortex; TP, temporal pole. a.u., arbitrary units; ms, milliseconds; pref., preferred; vu, virtual units. Error bars indicate SEM. \*\**P* < 0.01; \*\*\**P* < 0.001.

Direction cells were particularly present in parahippocampal cortex and hippocampus (Figure 4E). Across the population, their preferred directions were broadly distributed, but showed a bias towards a large mountain in the environment’s background (binomial test, P = 0.004; Figures 4F and 4G). This bias of preferred directions towards a prominent distal landmark resembles findings in rodents (Acharya et al., 2016) and provides neural evidence for the importance of visual cues in human spatial navigation (Ekstrom, 2015; Ekstrom et al., 2018), which may stabilize the transformation circuits (Bicanski and Burgess, 2018). Converging projections from anchor cells and direction cells may then lead to allocentric spatial representations in downstream neurons.

### Anchor cells support spatial memory

Beyond navigation, do anchor cells have a role in spatial memory? To answer this question, we examined (i) whether characteristics of “memory-sensitive cells” were common among anchor cells; (ii) whether the tuning strengths of anchor cells varied with spatial memory performance; and (iii) whether anchor cells showed conjunctive object tuning.

First, we examined whether characteristics of memory-sensitive cells, which changed their firing in relation to the patients’ spatial memory performance, were common among anchor cells. We found many memory-sensitive cells in various medial temporal lobe regions (Figure 5A), including neurons that increased (*n* = 59; binomial test, *P* < 0.001; e.g., Figure 5B) or decreased (*n* = 69; binomial test, *P* < 0.001; e.g., Figure 5C) their firing rates in association with good memory performance. The finding of both positive and negative memory-sensitive cells is in line with previous human single-neuron recordings (Tsitsiklis et al., 2020), and negative memory-sensitive cells may relate to inhibitory engrams, which are negative replicas of excitatory memory representations (Barron et al., 2017). We observed that anchor cells were strongly integrated into the memory network, as memory-related firing rate changes were common among anchor cells (29 of 95 anchor cells fulfilled the criteria for being memory-sensitive cells; 30.5%; chi-squared test, *χ*^2^ = 12.691, *P* < 0.001). In comparison, characteristics of memory-sensitive neurons were not particularly common among direction or place cells (23.8% and 21.1%, respectively; both *χ*^2^ < 2.563, both P > 0.109; Figure 5D).

**Figure 5.**
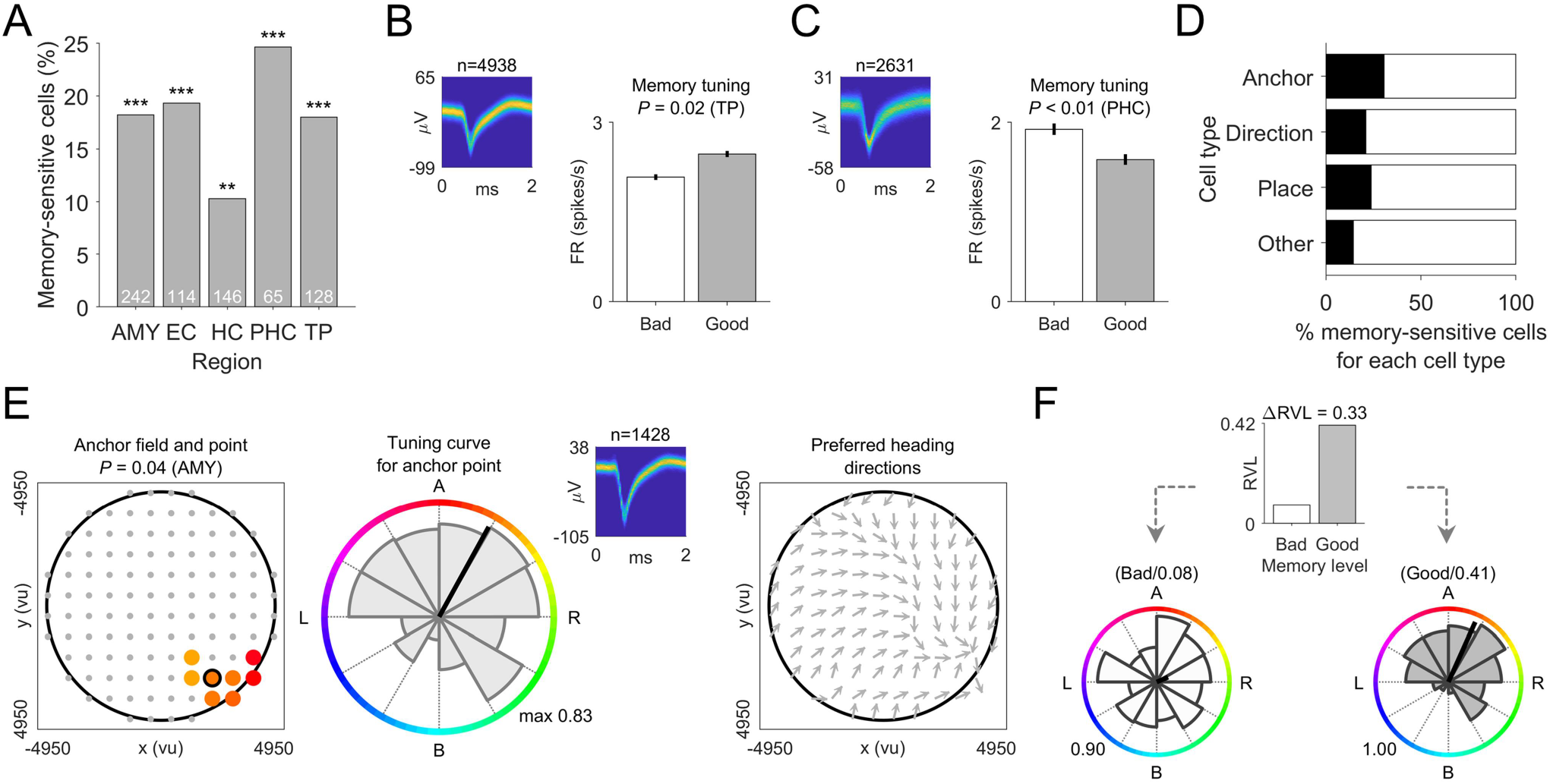
Anchor cells support spatial memory. (**A**) Distribution of memory-sensitive cells across brain regions. (**B and C**) Examples of memory-sensitive cells increasing (**B**) or decreasing (**C**) their firing rate in relation to good versus bad memory performance. (**D**) Prevalence of characteristics of memory-sensitive cells among anchor cells, direction cells, place cells, and other cells. (**E and F**) Example anchor cell (**E**) exhibiting more precise tuning during periods with better memory performance (**F**). A (B; L; R), anchor point ahead (behind; to the left; to the right) of the subject. AMY, amygdala; EC, entorhinal cortex; HC, hippocampus; PHC, parahippocampal cortex; TP, temporal pole. max, maximum firing rate; ms, milliseconds; RVL, Rayleigh vector length; vu, virtual units. Error bars indicate SEM. \*\**P* < 0.01; \*\*\**P* < 0.001.

Second, to understand whether anchor-cell properties varied with spatial memory performance, we examined the relationship between performance and the precision of the anchor cells’ directional tuning. We observed that a subset of anchor cells showed more precise anchor-cell tuning (i.e., higher Rayleigh vector lengths of the tuning curve) during task periods when subjects showed better spatial memory performance (11 of 95; binomial test, P = 0.008; e.g., Figures 5E and 5F). This phenomenon was more common among anchor cells that had preferred anchor bearings pointing “ahead” as compared to those with preferred anchor bearings pointing “behind” (chi-squared test, *χ*^2^ = 4.117, *P* = 0.042). Sharper tuning of anchor cells with anchor points ahead of the navigating organism may thus contribute to successful spatial navigation and accurate spatial memory.

Memory recall is often triggered by a sensory cue, followed by pattern-completion processes that reinstate the original spatial context associated with the cue (Staresina and Wimber, 2019). In a third step, we thus examined the relationship between anchor cells and object cells (*n* = 118; binomial test, *P* < 0.001; Figure 6), which provided neural representations of the objects whose locations had to be learned and retrieved throughout the task. For example, one object cell increased its activity during trials in which the patient had to remember and re-encode the location of a “globe” object (Figure 6A). Object cells represented mainly non-spatial information about the objects, because—when examining object cells that exhibited more than one preferred object—the locations of the preferred objects were not spatially closer to each other as compared to similar numbers of randomly selected object locations (permutation test, P = 0.464). We found that a significant number of anchor cells fulfilled the criteria for being an object cell (27 of 95; binomial test, *P* < 0.001; chi-squared test, = 12.052, *P* < 0.001; Figure 6F). These conjunctive anchor–object cells may constitute a neural interface between spatial and non-spatial task features, which could be instrumental in reinstating the egocentric spatial context upon a non-spatial cue during memory recall.

**Figure 6.**
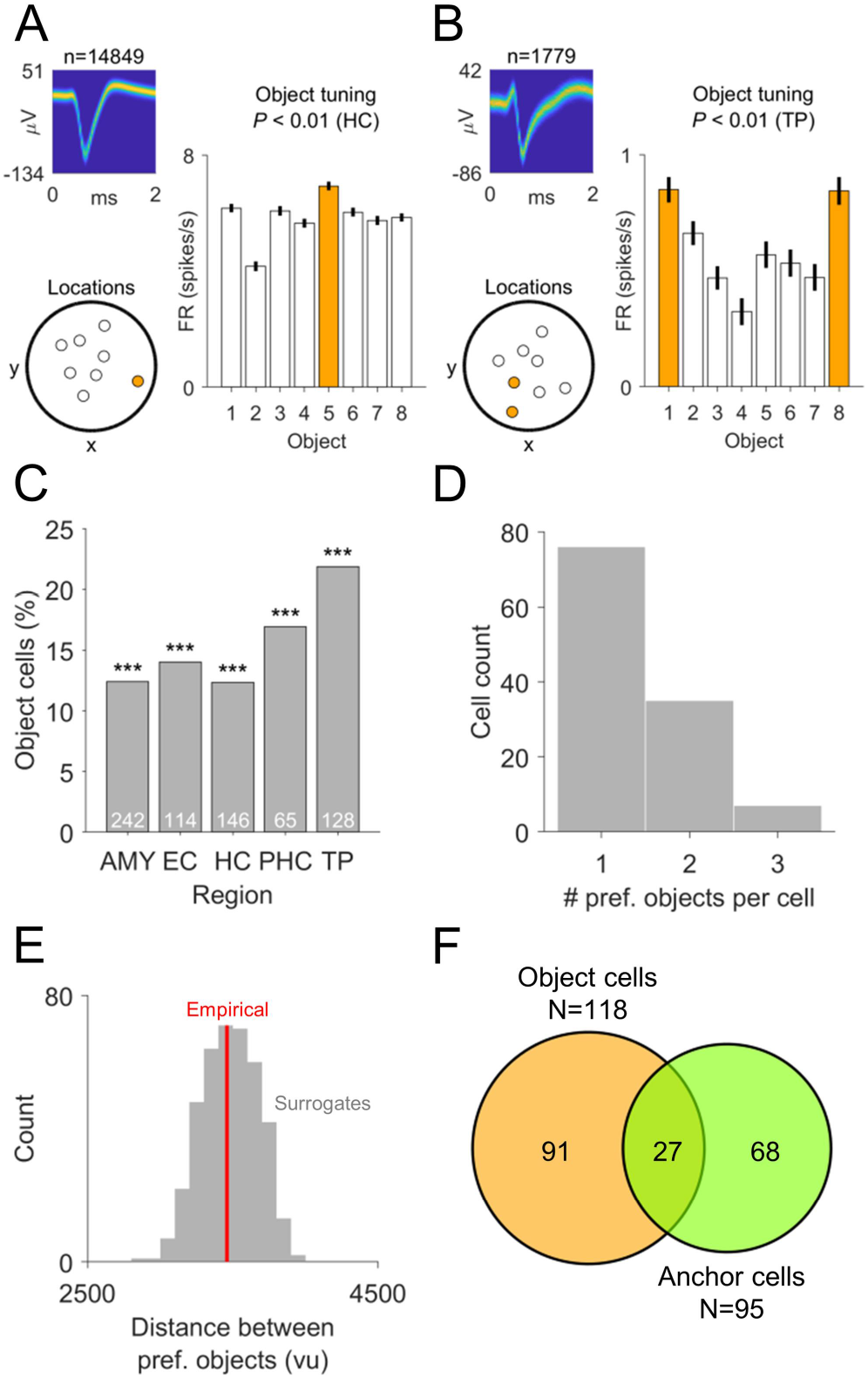
Relationship of anchor cells to object cells whose locations have to be learned and retrieved throughout the task. (**A and B**) Examples of object cells. Each bar shows the average firing rate during trials with a given object. Orange bars depict the cells’ preferred object(s). For example, the object cell shown in (**A**) activated during trials in which object #5 (in this case, the globe) had to be retrieved and re-encoded. Upper left subpanels show spikedensity plots (number above subpanel indicates spike count). Lower left subpanels show the object locations (locations of preferred objects are indicated in orange). (**C**) Distribution of object cells across brain regions. (**D**) Histogram of the number of preferred objects per object cell. (**E**) Average distance between preferred objects for object cells with at least two preferred objects. Red line, empirical average distance between preferred objects; gray bars, histogram of surrogate distances. (**F**) Overlap between object cells and anchor cells. AMY, amygdala; EC, entorhinal cortex; HC, hippocampus; PHC, parahippocampal cortex; TP, temporal pole. FR, firing rate; pref, preferred. Error bars indicate SEM. \*\*\**P* < 0.001.

## Discussion

Humans encode, store, and recall information about places, distances, and directions in both allocentric and egocentric reference frames (Burgess, 2006). Here, using single-neuron recordings in neurosurgical epilepsy patients performing a virtual navigation task, we identified anchor cells whose activity encoded the subject’s egocentric direction towards local reference points in the virtual environment. Anchor cells supported full vectorial representations of egocentric space by additionally encoding the distances to anchor points. Anchor cells were abundant in parahippocampal cortex and were integrated into a neural memory network. Our identification of anchor cells contributes to the mechanistic understanding of how the human brain supports spatial navigation and spatial memory (Table S2). Specifically, our data suggest that anchor cells constitute an egocentric counterpart of place cells and cells encoding allocentric direction, thus providing a neural code for egocentric spatial maps.

### The relationship between anchor cells and other egocentric spatial representations

Previous studies identified neurons in animals whose firing rates encoded the egocentric spatial relationship between the navigating animal and salient features of the environment, such as the environment’s center, its boundaries, or objects placed within the environment (Alexander et al., 2020; Gofman et al., 2019; Hinman et al., 2019; LaChance et al., 2019; Sarel et al., 2017; Wang et al., 2018; Wilber et al., 2014).

In contrast, anchor cells had anchor points in many different parts of the spatial environment. Anchor cells may therefore support a comprehensive cognitive map of the spatial environment, whereas the previously discovered egocentric spatial representations may provide egocentric information about specific aspects of the environment. The distinction between anchor cells and neurons that are egocentrically tuned to salient environmental features may thus be analogous to the relationship between place cells and neurons that are allocentrically tuned to such environmental features: whereas place cells have place fields in various locations of the environment (Duvelle et al., 2019; Gauthier and Tank, 2018), object-vector cells and boundary-vector cells activate when environmental objects and boundaries are at specific allocentric directions and distances from the animal (Deshmukh and Knierim, 2013; Høydal et al., 2019; Lever et al., 2009).

Alternatively, one may hypothesize that anchor cells constitute a generalized version of other egocentric spatial cell types because—depending on task demands—anchor cells may shift their anchor points towards the locations of external items, the environment center, or goal locations. Such shifts could result in cellular activity as described for egocentric cue direction cells (Wilber et al., 2014), item-bearing cells (Wang et al., 2018), center-bearing cells (LaChance et al., 2019), or goal-vector cells (Sarel et al., 2017). In our study, we may not have seen clear shifts of anchor points towards object locations due to the fact that the participants had to learn and retrieve the locations of multiple different objects—potentially forcing the participants to create egocentric spatial maps that represented the different parts of the spatial layout similarly strong. Hence, future studies are needed to elucidate the mechanisms by which anchor points arise and whether anchor points are allocated depending on task demands.

It is notable though that a number of anchor cells exhibited anchor points in the center of the environment—underscoring the idea that the geometrical center of an environment has a distinct role for orientation and navigation (Gallistel, 1990; LaChance et al., 2019). These anchor cells with center anchor points may be related to the previously observed center-bearing cells in the rat postrhinal cortex (LaChance et al., 2019) and to path cells described in the human entorhinal cortex, which activate when epilepsy patients navigate clockwise or counterclockwise on a square road in a virtual environment (Jacobs et al., 2010).

### Anchor cells’ role in the processing hierarchy of the brain’s spatial navigation system

We hypothesize that anchor-cell functioning constitutes an essential component in the processing hierarchy of the brain’s spatial navigation system. Our results suggest that anchor cells extract egocentric spatial information from sensory input during navigation. Then, as suggested by theoretical models (Bicanski and Burgess, 2018; Wang et al., 2020), neural transformation circuits may combine anchor-cell activity with signals from cells encoding allocentric direction such as head-direction cells (Taube et al., 1990). Integrating anchor-cell activity with direction-cell activity would allow the transformation of anchor-cell activity into allocentric coordinates, by means of which neural activity could emerge as described for spatial view cells (Rolls, 1999), spatial target cells (Tsitsiklis et al., 2020), or place cells that represent remote locations (Jercog et al., 2019; Pfeiffer and Foster, 2013; Shahi et al., 2019).

We assume that the transformation of anchor-cell activity into allocentric coordinates proceeds in parallel to the transformation of activity from item-bearing cells and egocentric boundary cells (Alexander et al., 2020; Gofman et al., 2019; Hinman et al., 2019; Wang et al., 2018; Wilber et al., 2014) into allocentric neuronal activity as seen in object-vector cells and boundary-vector cells (Deshmukh and Knierim, 2013; Høydal et al., 2019; Lever et al., 2009), respectively. These transformations of egocentric into allocentric spatial codes are probably useful for enabling long-term storage and abstract, self-independent knowledge (Bellmund et al., 2018; Burgess, 2006; Buzsáki and Moser, 2013; Epstein et al., 2017).

To enable movement planning and movement execution in service of navigation, allocentric spatial representations may be transformed back into egocentric spatial representations in order to determine action plans from the first-person perspective (such as “navigate forward and then to the left”). We thus propose that anchor cells also play a role in route planning, movement planning, and movement execution. To this end, sequences of anchor-cell activity may be forwarded to motor cortices to transform them into directly executable body movements (Olson et al., 2020). To provide anatomical evidence for this suggestion, future studies should examine the existence of anchor cells in (supplementary) motor cortices and how they relate to anchor cells in medial temporal lobe regions. Taken together, we assume that anchor cells participate in both the ascending and descending part of a neural circuit for orientation and navigation, assuring that the navigating organism is accurately oriented in its spatial environment and able to successfully execute navigation routes through the environment.

### Anchor cells and the parahippocampal cortex

We found that anchor cells were particularly prevalent in the parahippocampal cortex. This finding parallels the existence of other egocentrically tuned spatial cell types observed in the rat postrhinal cortex (Gofman et al., 2019; LaChance et al., 2019). The parahippocampal cortex is located at the junction between visual cortices and the hippocampal formation (Aminoff et al., 2013), lending anatomical support to the hypothesis that anchor cells help to transform percepts into allocentric spatial representations. However, an open question is whether anchor-cell activity originates in the parahippocampal cortex or whether the parahippocampal cortex inherits anchor-cell activity from other brain regions such as the parietal cortex, which has long been implicated in egocentric spatial processing (Aguirre and D’Esposito, 1999; Burgess, 2008). Solving this question will help to anatomically pinpoint the processing hierarchy of the brain’s spatial navigation system.

Previous studies in humans using functional magnetic resonance imaging, intracranial electroencephalography, or single-neuron recordings revealed that a subpart of the parahippocampal cortex reliably responds to images of spatial scenes (Bastin et al., 2013; Epstein and Kanwisher, 1998; Mormann et al., 2017). This subpart has thus been termed “parahippocampal place area” (Epstein and Kanwisher, 1998). The abundance of anchor cells and other egocentric spatial cell types in the parahippocampal cortex could provide a mechanistic explanation for the neural responses of the parahippocampal place area to spatial scenes, because the presentation of spatial scenes from a first-person perspective may cause increased activity among anchor cells, center-bearing cells, and egocentric boundary cells as a subject attempts to orient within the viewed spatial scene.

Further, we suggest that the high prevalence of anchor cells in the parahippocampal cortex could be relevant clinically by explaining why parahippocampal lesions cause disruptions of performance on spatial navigation and memory tasks that require egocentric reference frames [e.g., (Ploner et al., 2000; Weniger and Irle, 2006)]. A reduced number of anchor cells due to the removal of the parahippocampal cortex may have accounted for impaired navigation and memory in these patients.

### Possible functions for anchor cells in navigation and memory

The activity of anchor cells provides the subject with a sense of how the proximal spatial layout is oriented relative to the subject’s viewpoint. Across the population of anchor cells, anchor points are positioned in many different locations of the surrounding environment, providing the subject with a comprehensive egocentric mental map of the local environment’s extent and shape. Anchor cells may thus constitute a neural substrate for egocentric navigation strategies in human spatial behavior (Burgess, 2006; Coughlan et al., 2018; Ekstrom and Isham, 2017; Ekstrom et al., 2018; Wang and Spelke, 2002).

In addition to providing orientation, anchor cells may allow route planning from a first-person perspective: Via sequential activation of anchor cells with neighboring anchor points, navigation paths may be simulated from an egocentric viewpoint in order to plan future routes that start from the subject’s current location and orientation. Such preparatory activity in service of goal-directed navigation has been observed in hippocampal place cells (Pfeiffer and Foster, 2013), but has yet to be demonstrated for egocentric spatial representations. Thus, more broadly, anchor-cell activity may be used for imagining future scenarios from egocentric viewpoints (Schacter et al., 2007).

In the memory domain, the distinction between allocentric and egocentric spatial representations is thought to be paralleled by the contrast between semantic and episodic memories (Buzsáki and Moser, 2013): Whereas semantic memories consist of knowledge about the world that is abstracted from personal experiences, episodic memories are past experiences remembered from the first-person perspective (Binder and Desai, 2011). Given that anchor cells provide a neural substrate for egocentric spatial representations, they may therefore contribute to the neural basis of episodic memories. Our findings already indicate that anchor cells have a role in spatial memory beyond pure navigation, because anchor cells’ firing rates and directional tuning strengths correlated with the subject’s spatial memory performance (Figure 5). Moreover, we found that a number of anchor cells showed characteristics of object cells by increasing their firing rates during trials when the locations of one or more specific objects had to be retrieved and re-encoded. Such conjunctive anchor–object cells provide an interface between spatial and non-spatial task features and enable, in principle, that a non-spatial memory cue triggers the recall of the egocentric spatial context of the associated memory.

### Conclusion

In this study, we identified a neural code for egocentric spatial maps in the human medial temporal lobe. Anchor cells appeared to constitute this code’s key unit by encoding egocentric directions between local reference points in the spatial environment and the navigating subject. Anchor cells may thus provide the subject with an egocentric representation of its proximal environment, thereby allowing the use of egocentric navigation strategies. Possible functions of anchor cells extend to route planning and the imagination of future scenarios from egocentric viewpoints. Anchor cells may finally underlie the first-person perspective in episodic memories to enable a vivid recollection of past experiences.

## Acknowledgements

We are grateful for all the patients who volunteered to participate in this study. We thank the clinical team of the Freiburg Epilepsy Center, Germany, for their continuous support, D. Lachner-Piza for technical assistance, and S. Qasim, M. Peer, and D. Schonhaut for comments on the manuscript.

L.K., A.B., E.T.R., R.K., and A.S.-B. were supported by the Federal Ministry of Education and Research (BMBF; 01GQ1705A). L.K., A.B., M.T., A.S.-B., and J.J. received funding via National Science Foundation (NSF) grant BCS-1724243. L.K., A.B., and A.S.-B. were supported by National Institutes of Health (NIH) grant 563386 and NIH/NINDS grant U01 NS1113198-01. L.K. was supported by a travel grant from the Boehringer Ingelheim Fonds (Mainz, Germany). P.C.R. received research grants from the Fraunhofer Society (Munich, Germany) and from the Else Kröner-Fresenius Foundation (Bad Homburg, Germany) and personal fees, travel support, and honoraria for lectures from Boston Scientific (Marlborough, MA, USA), Brainlab (Munich, Germany) and Inomed (Emmendingen, Germany). N.A.H. received funding via the German Research Foundation (DFG; HE 8302/1-1). B.P.S. was supported by a Wellcome Trust/Royal Society Sir Henry Dale Fellowship (107672/Z/15/Z). M.T. and J.J. were supported by NIH grant MH104606. M.J.K. was supported by NIH grants MH55687 and MH061975.

## Author contributions

L.K. and J.J. designed the study; L.K., A.B., P.C.R, and A.S.-B. recruited participants; P.C.R. implanted electrodes; L.K. and A.B. collected data; L.K. performed analyses with contributions from B.P.S., E.T.R., C.T.W., N.A.H., M.T., and R.K.; A.S.-B., J.J., M.J.K., and R.K. acquired the necessary resources and financial support as well as managed and supervised the project; L.K. and J.J. wrote the paper; all authors reviewed and edited the final manuscript.

## STAR Methods

### Experimental model and subject details

14 human subjects (seven female; age range, 19-51 years; mean age ± SEM, 33.1 ± 3.0 years) undergoing treatment for pharmacologically intractable epilepsy participated in the study. Informed written consent was obtained from all patients. The study conformed to the guidelines of the ethics committee of the University Hospital Freiburg, Freiburg im Breisgau, Germany.

### Neurophysiological recordings

Patients were surgically implanted with intracranial depth electrodes in the medial temporal lobe for diagnostic purposes in order to isolate the epileptic seizure focus for potential subsequent surgical resection. The exact electrode numbers and locations varied across subjects and were determined solely by clinical needs. Neuronal signals were recorded using Behnke-Fried depth electrodes (Ad-Tech Medical Instrument Corp., Racine, WI). Each depth electrode contained a bundle of nine platinum-iridium microelectrodes with a diameter of 40 μm that protruded from the tip of the depth electrode (Fried et al., 1999). The first eight microelectrodes were used to record action potentials and local field potentials. The ninth microelectrode served as reference. Microelectrode coverage included amygdala, entorhinal cortex, fusiform gyrus, hippocampus, parahippocampal cortex, temporal pole, and visual cortex. We recorded microwire data at 30 kHz using NeuroPort (Blackrock Microsystems, Salt Lake City, UT).

### Spike detection and sorting

Neuronal spikes were detected and sorted using Wave_Clus (Chaure et al., 2018). We used default settings with the following exceptions: “template_sdnum” was set to 1.5 to assign unsorted spikes to clusters in a more conservative manner; “min_clus” was set to 60 and “max_clus” was set to 10 in order to avoid over-clustering; and “mintemp” was set to 0.05 to avoid under-clustering. All clusters were visually inspected and judged based on the spike shape and its variance, inter-spike interval (ISI) distribution, and the presence of a plausible refractory period [following (Kutter et al., 2018)]. If necessary, clusters were manually adjusted or excluded. Furthermore, clusters were excluded that exhibited mean firing rates of <0.1 Hz during the analysis time window [following (Ekstrom et al., 2003)]. Spike waveforms are shown as spike-density plots in all figures [following (Reber et al., 2019)].

In total, we identified *N* = 729 clusters (also referred to as “neurons” or “cells” throughout the manuscript) across 18 experimental sessions from all 14 patients. Because different sessions from the same patient were separated by >12 h, we treated neuronal responses of these sessions as statistically independent units. An experienced rater (B.P.S.) assigned the tips of depth electrodes to brain regions based on post-implantation MRI scans in native space so that neurons recorded from the corresponding microelectrodes could be assigned to these regions. We recorded *n* = 242 neurons from amygdala, *n* =114 neurons from entorhinal cortex, n = 25 neurons from fusiform gyrus, n = 146 neurons from hippocampus, n = 65 neurons from parahippocampal cortex, n = 128 neurons from temporal pole, and n = 9 from visual cortex. Due to low numbers of neurons in fusiform gyrus and visual cortex, we excluded these regions from region-specific analyses.

For recording quality assessment (Figure S2), we calculated the number of units recorded on each wire; the ISI refractoriness for each unit; the mean firing rate for each unit; and the waveform peak signal-to-noise ratio (SNR) for each unit. The ISI refractoriness was assessed as the percentage of ISIs with a duration of <3 ms. The waveform peak SNR was determined as: SNR = A_peak_/STD_noise_, where A_peak_ is the absolute amplitude of the peak of the mean waveform, and STD_noise_ is the standard deviation of the raw data trace (filtered between 300 and 3,000 Hz).

### Task

During experimental sessions, patients sat in bed and performed an object–location memory task on a laptop computer (Figure 1), which was adapted from previous studies (Chen et al., 2018a; Doeller et al., 2008, 2010; Kunz et al., 2015, 2019b).

During the task, patients first learned the locations of eight everyday objects by collecting each object from its location once (this initial learning phase was excluded from all analyses). Afterwards, patients completed variable numbers of test trials (Figure 1A) depending on compliance. Each test trial started with an inter-trial-interval of 3-5 s duration (uniformly distributed). Patients were then presented with one of the eight objects (“cue”; duration of 2 s). During the subsequent retrieval period (“retrieval”; self-paced), patients navigated to the assumed object location and indicated their arrival via a button press. Next, patients received feedback on the accuracy of their response using one of five different emoticons (“feedback”; duration of 1.5 s). Response accuracy was measured as the Euclidean distance between the response location and the correct location (“drop error”). The retrieved object then appeared in its correct location and patients collected it from there in order to further improve their associative object-location memories (“re-encoding”; self-paced).

Drop errors were transformed into spatial memory performance values by ranking each drop error within 1 million potential drop errors. Potential drop errors were the distances between the trial-specific correct object location and random locations within the virtual environment. This transformation accounted for the fact that the possible range of drop errors is smaller for object locations in the center of the virtual environment as compared to object locations in the periphery of the virtual environment (Miller et al., 2018). A spatial memory performance value of 1 represents the smallest possible drop error, whereas a spatial memory performance value of 0 represents the largest possible drop error.

The virtual environment comprised a grassy plain [diameter of 10,000 virtual units (vu)] surrounded by a cylindrical cliff. There were no landmarks within the environment. The background scenery comprised a large and a small mountain, clouds, and the sun (Figure 1A and 1B). All distal landmarks were rendered at infinity and remained stationary throughout the task.

Patients navigated the virtual environment using the arrow keys of the laptop computer (forward, turn left, turn right). Instantaneous virtual locations and heading directions (which are identical with viewing directions in our task) were sampled at 50 Hz. We aligned the behavioral data with the electrophysiological data via visual triggers, which were detected by a phototransistor attached to the screen of the laptop computer. The phototransistor signal was recorded together with the electrophysiological data at a temporal resolution of 30 kHz.

### General statistical procedure

At each time point (sampled at 50 Hz), the participant’s allocentric direction and location was given by the yaw value and the (x/y)-coordinate of the virtual character’s position in the virtual environment, respectively. Neuronal spike times were adjusted to the behavioral time axis according to the trigger time stamps. We then downsampled the behavioral data to 10 Hz [following (Jacobs et al., 2010)] and calculated the neuronal firing rate (spikes/s) for each time bin. Time periods in which the patient remained stationary for >2 s were excluded from the analysis.

To identify different cell types, we employed an ANOVA framework (Ekstrom et al., 2003; Manns et al., 2007; Qasim et al., 2019; Tsitsiklis et al., 2020; Wood et al., 2000), in which we assessed the effects of different factors on firing rates. To identify direction and place cells, we used a two-way ANOVA with factors “direction” and “place”. To identify anchor cells, we used a three-way ANOVA with factors “direction”, “place”, and “anchor bearing”. To identify object cells, we used a three-way ANOVA with factors “direction”, “place”, and “object”. In all ANOVAs (computed via MATLAB’s *anovan* function), we used Type II sums of squares, which controls for main effects of other factors when determining significance of a given factor. Empirical *F*-values of a given factor were considered significant, when they exceeded the 95^th^ percentile of 101 surrogate *F*-values which we obtained by performing the same ANOVA on circularly shifted firing rates [with the end of the session wrapped to the beginning; following, e.g., (Qasim et al., 2019)].

Tuning curves are displayed as the estimated marginal means of a given factor when controlling for the other factors (computed via MATLAB’s *multcompare* function), inspired by analysis procedures in rodents that identify independent effects of different factors on firing rates (Burgess et al., 2005; Hardcastle et al., 2017).

All analyses were carried out in MATLAB 2018b using MATLAB toolboxes and custom MATLAB scripts. Unless otherwise indicated, we considered results statistically significant when the corresponding *P*-value fell below an alpha-level of α = 0.05. Analyses were twosided, if not otherwise notified. Binomial tests evaluated the significance of proportions of neurons relative to a chance level of 5% (two-sided), if not otherwise specified. Surrogate statistics were generally one-sided to assess whether an empirical test statistic exceeded a distribution of surrogate statistics significantly (Oostenveld et al., 2011). Statistics on angular data were carried out using the CircStat toolbox (Berens, 2009). The significance of overlaps between different cell types was assessed using chi-squared tests.

### Anchor cells

We identified anchor cells using a two-step procedure (Figure 1F and 1G). In the first step, separately for each candidate anchor point, we analyzed each neuron’s firing rate via a threeway ANOVA with factors “direction”, “place”, and “anchor bearing” to assess the relevance of “anchor bearing” while controlling for “direction” and “place”. We calculated anchor bearings as the angular difference between the subject’s instantaneous heading angle and the concurrent angle of the vector from the subject’s location to the anchor point (Figure 1E). Candidate anchor points (n = 112) were evenly distributed across the virtual environment (distance between neighboring candidate anchor points, 900 vu; Figure 1F). No candidate anchor points were located outside the circular boundary. The factors “direction” and “anchor bearing” could take on one of twelve values (angular resolution, 30°). The factor “place” could take on one of 100 values representing a 10 x 10 grid overlaid onto the virtual environment (bin edge length, 900 vu). Only factor levels with ≥5 separate observations (for example, five temporally distinct visits to location bin *i*) were included to ensure sufficient behavioral sampling. For the factor “anchor bearing” we then extracted the raw ANOVA *F*-value (*F*_empirical_) and the corresponding estimated firing rate map (eFR_empirical_), which is the tuning curve of the firing rate as a function of “anchor bearing” while controlling for “direction” and “place” (for examples, see the middle column of Figure 2). Using the circular-shift procedure described above, we estimated surrogate *F*-values (*F*_surrogate_) and surrogate estimated firing rate maps (eFR_surrogate_). A candidate anchor point was considered significant, if its *F*_empirical_ value exceeded the 95^th^ percentile of its *F*_surrogate_ values (corresponding to *P* < 0.05).

In the second step, we employed cluster-based permutation testing (Oostenveld et al., 2011) to assess the overall significance of the cell regarding anchor-bearing tuning: contiguous clusters of significant candidate anchor points were identified and their percentiles of *F*_empirical_ within Fsurrogate were summed up, resulting in a cluster-percentileempirical value (for example, a contiguous cluster of 10 significant candidate anchor points, where all candidate anchor points had a percentile value of 97%, resulted in a cluster-percentile value of 970%). We considered this cluster-percentile_empirical_ value statistically significant, if it exceeded the 95^th^ percentile of surrogate cluster-percentile_surrogate_ values (corresponding to *P* < 0.05). Here, cluster-percentile_surrogate_ values were created by using each of the *F*_surrogate_ matrices as a hypothetical empirical matrix once, each time assessing its cluster-percentile_surrogate_ value by comparing it against all other matrices (both the remaining *F*_surrogate_ matrices and the *F*_empirical_ matrix), as described above.

For each anchor cell, we show the contiguous cluster of significant candidate anchor points (i.e., the “anchor field”; e.g., Figure 2A, left): each significant candidate anchor point is depicted as a colored, bold dot; non-significant candidate anchor points are indicated as gray, small dots. Coloring corresponds to the circular mean of the estimated firing rate map eFRempirical for that candidate anchor point (for example, red means that the neuron’s firing rate increased when the subject was moving towards this point; cyan means that the neuron’s firing rate increased when the subject was moving away from this point). We obtained each cell’s “anchor point” by calculating the center of mass of the anchor field using MATLAB’s *regionprops* function. We note that the terms “anchor field” and “anchor point” have been used before to describe the tuning of hippocampal place cells towards reference points (Shahi et al., 2019).

#### Vector-field maps

As an approximate illustration of the cell’s anchor-bearing tuning, we additionally show the cell’s preferred allocentric direction as a function of location (e.g., Figure 2A, right). Here, the location-specific allocentric direction tuning curve is estimated via a two-way ANOVA with factors “direction” and “place”, which takes only data points into account when the subject is in the vicinity of the corresponding candidate anchor point [the vicinity of a candidate anchor point is defined as the point’s (x/y)-coordinate ± 3,333 vu]. For example, the vector-field map in Figure 2A, right, shows that allocentric direction tuning of this cell varies across different locations, twisting towards a spot in the northeast part of the virtual environment. This vector-field map can thus illustrate the anchor-cell plot. Of note, the vector-field map does not match the anchor-cell plot closely in cases when direction and anchor bearing explain relevant and independent amounts of variance in the firing rates. This is due to the fact that the anchor-cell plot shows anchor-bearing tuning while accounting for the effects of direction and location (three-way ANOVA with factors “direction”, “place”, and “anchor bearing”), whereas the vector-field map shows direction tuning while only accounting for the effect of location (two-way ANOVA with factors “direction” and “place”).

#### Preferred bearing of anchor cells

For each anchor cell, we extracted its preferred bearing towards the anchor point via the circular mean of the corresponding tuning curve. Bimodality of preferred anchor bearings was tested by applying a Rayleigh test on the preferred anchor bearings multiplied by two.

#### Spatial organization of anchor points

We evaluated bimodality of anchor-point distances towards the environmental center using Hartigan’s dip test (Hartigan and Hartigan, 1985) and separated the two groups using MATLAB’s *kmeans* function.

#### Distance tuning of anchor cells

To investigate whether anchor cells encoded the distance towards the anchor point, we analyzed each anchor cell’s firing rate as a function of Euclidean distance towards the anchor point. Only time points, when the subject’s current anchor bearing was in alignment with the cell’s preferred anchor bearing (±90°) were considered for this analysis. We then tested whether firing rates significantly increased or decreased when the subject was “far” from or “close” to the anchor point (median split of distance values). To this end, we performed a two-sample *t*-test between firing rates associated with far versus close distances from the anchor point and extracted the resulting *t*-statistic (*t*_empirical_) for each anchor cell. To obtain surrogate *t*-statistics (*t*_surrogate_), we circularly shifted the empirical firing rates 401 times in relation to the behavioral data. We considered a cell to exhibit distance tuning, if *t*empirical exceeded the 95^th^ percentile of *t*_surrogate_ values (positive distance tuning) or if it fell below the 5^th^ percentile of *t*_surrogate_ values (negative distance tuning).

#### Anchor-cell tuning and memory performance

To establish a relationship between the specificity of the tuning curve of anchor cells and spatial memory performance, we divided the data into two data parts based on a median split of spatial memory performance values (resulting in two memory levels: “good” and “bad”). For each data part, we used the three-way ANOVA with factors “direction”, “place”, and “anchor bearing” towards the anchor point (identified using the entire data) to extract the estimated firing rate map eFR_empirical-part_ for the factor “anchor bearing”. We estimated the precision of eFR_empirical-part_ as the Rayleigh vector length (RVL), separately for each data part (RVL_part_), and calculated the difference between data part-specific RVLs as: ΔRVL = diff(RVL_part_). For each anchor cell, we then tested whether its ΔRVL value was above the 95^th^ percentile of 401 surrogate ΔRVL values obtained by performing the analysis on circularly shifted firing rates. We tested whether the proportion of these memory-modulated anchor cells were more common among anchor cells with preferred anchor bearings pointing “ahead” (i.e., “ahead” ± 90°) as compared to anchor cells with preferred anchor bearings pointing “behind” (i.e., “behind” ± 90°) using a chi-squared test.

### Direction cells and place cells

Rodent head-direction cells (Taube et al., 1990) activate whenever an animal’s head is pointing in a specific global direction that is defined relative to a world-referenced coordinate system (for example, when the head is pointing “north” or “south”). Here, we identified “direction cells” that exhibited firing-rate modulations as a function of the patients’ current heading direction within the virtual environment.

To identify direction cells, we analyzed each neuron’s activity by means of a two-way ANOVA with factors “direction” and “place”. The factor “direction” could take on one of twelve values (angular resolution, 30°). The factor “place” could take on one of 100 values representing a 10 x 10 grid overlaid onto the virtual environment (bin edge length, 900 vu). Only factor levels with ≥5 separate observations were included to ensure sufficient behavioral sampling. We then extracted the raw ANOVA *F*-value for the factor “direction” (*F*_empirical_) and the estimated firing rate map (eFR_empirical_), while controlling for the factor “place”. We calculated statistical significance of *F*_empirical_ values using surrogate statistics as described above. For each direction cell, we extracted its preferred direction via the circular mean of the directional tuning curve.

To test whether preferred directions were biased towards the distal landmarks, we divided the background scenery into the following parts: large mountain, large gap, small mountain, and small gap. For each landmark we then counted how often a preferred direction pointed towards it (*N*_observed_). We tested for significance of landmark-specific *N*_observed_ values using binomial tests against chance levels, which were separately estimated for each landmark based on its angular extension.

To compare the vector-field maps of direction cells with the vector-field maps of anchor cells, we computed “vector-field strengths” as the Rayleigh vector length of all vectors in the vector-field map (a completely homogeneous vector-field map would result in a Rayleigh vector length of 1.0; a completely inhomogeneous vector-field map would result in a Rayleigh vector length of 0.0).

We identified place cells via the same procedure as described for direction cells using a two-way ANOVA with factors “direction” and “place”. We defined place bins as those spatial bins in which the empirical firing rate exceeded the 95^th^ percentile of surrogate firing rates [following (Ekstrom et al., 2003)].

### Memory-sensitive cells

We identified memory-sensitive cells as those cells exhibiting significantly increased or decreased firing rates during periods with good versus bad spatial memory performance (median split of spatial memory performance values). To this end, we performed a two-sample *t*-test for each cell between firing rates associated with good versus bad spatial memory performance and extracted the resulting *t*-statistic (*t*_empirical_). Surrogate statistics were obtained as described above. We labeled a neuron as “memory-sensitive”, if *t*_empirical_ exceeded the 95^th^ percentile of *t*_surrogate_ values (positive memory-sensitive cell) or if it fell below the 5^th^ percentile of *t*_surrogate_ values (negative memory-sensitive cell).

### Object cells

To identify object cells, we analyzed each neuron’s activity using a three-way ANOVA with factors “direction”, “place”, and “object”. The factor “object” could take on one of eight different values (because each patient learned and retrieved the locations of eight different objects). For all time bins of a given trial, the factor “object” had the same value. We obtained object cells and preferred objects [i.e., objects for which the cell’s empirical firing rate exceeded the 95^th^ percentile of surrogate firing rates (Ekstrom et al., 2003)] using surrogate statistics as described above. Cells without a preferred object were excluded from the object-cell population.

To examine whether the preferred objects of object cells with ≥2 preferred objects exhibited a specific spatial relationship, we estimated the average Euclidean distance between the locations of all preferred objects, separately for each object cell with ≥2 preferred objects, and averaged across cells afterwards (*D*_empirical_). We created 401 surrogate values (*D*_surrogate_) for comparison by randomly selecting *n* object locations per cell, where *n* corresponds to the number of preferred objects of a given cell. We then determined the percentile of *D*_empirical_ within *D*_surrogate_ to test whether *D*_empirical_ was smaller than the 5^th^ percentile of *D*_surrogate_-values (in this case, locations of preferred objects would be closer to each other as expected by chance).

### Data and code availability

All data and code used to analyze the data are available upon request from the corresponding author. Data and code will be made publicly available after publication.

## Supplemental Information

### Supplemental Text

#### Text S1. Tuning strengths of spatially modulated neurons

Tuning strengths of spatially modulated cells in our study were generally lower than in rodents. For example, head-direction cells in rodents often exhibit baseline firing rates of about 0 spikes/s and increase their firing rates up to about 100 spikes/s at the preferred head direction (Taube et al., 1990). Directionally sensitive neurons in our study showed only moderate firing rate increases when subjects moved in the preferred direction [for similar tuning strengths, see for example (Ekstrom et al., 2003; Miller et al., 2013; Qasim et al., 2019; Tsitsiklis et al., 2020)]. The use of the classical terminology (e.g., the term “place cells” to refer to place-selective neurons) should thus be treated with caution (we adhered to these terms for ease of reading).

Different factors may contribute to the reduced selectivity of neuronal responses: In humans, it is not possible to adjust the localization of microelectrodes after implantation and a search for strongly tuned cells is thus not possible. Moreover, patients did not physically navigate the spatial environment, but rather completed a virtual navigation task, potentially associated with broader spatial tuning (Chen et al., 2018b). Additionally, neuronal firing in the human brain may be higher-dimensional than in rodents meaning that more internal and external factors (including ongoing thoughts, spontaneous occurrence of memories and ideas, and stimuli in the patient’s room) influence neuronal firing rates. Finally, (subtle) epileptogenic processes may have also affected the sharpness of the neurons’ tuning curves (Shuman et al., 2020). Reassuringly, control analyses showed a significant proportion of anchor cells after excluding neurons from the anchor-cell analysis that were recorded on microelectrodes located in brain regions potentially involved in the generation of epileptic seizures.

#### Text S2. Relationship between anchor cells and direction cells

Anchor cells encode egocentric direction towards local reference points, whereas direction cells encode allocentric direction. However, egocentric direction towards a reference point becomes increasingly similar to allocentric direction with increasing distance of the reference point from the subject.

In the main text, we showed that the homogeneity of vector-field maps differed between anchor cells and direction cells. We performed additional analyses to clarify the relationship between anchor cells and direction cells. First, to provide evidence that anchor-cell tuning did not spuriously arise due to potential collinearities between the factors “direction” and “anchor bearing” in our three-way ANOVA framework designed to identify anchor cells, we performed this ANOVA on surrogate data instead of empirical data [testing for significance of the tuning curves by comparing against other surrogate data; following (Kutter et al., 2018)]. In this way, we empirically estimated the percentage of anchor cells that may have arisen due to chance (for example, due to interdependencies between the factors “direction” and “anchor bearing”). As expected, this approach resulted in n = 34 (4.7%) statistically significant outcomes (i.e., false positives), confirming the *a priori* chosen alpha level of α = 0.05.

Second, we performed the anchor-cell analysis using a two-way ANOVA, with factors “place” and “anchor bearing”, in order to test for the number of anchor cells when not controlling for allocentric direction in our ANOVA framework. Using this approach, we observed n = 110 anchor cells (15.1%; binomial test, *P* < 0.001; as reported in the main text, we identified 95 anchor cells when also controlling for allocentric direction). Furthermore, anchor cell test statistics (i.e., each cell’s cluster-percentile_empirical_ value) were highly similar between both types of ANOVA (Spearman correlation, ρ_727_ = 0.626, *P* < 0.001) and the overlap between anchor cells identified via the two different analyses was significantly higher than expected by chance (chi-squared test, *χ*^2^ = 148.630, *P* < 0.001).

In sum, these analyses show (i) that anchor cells exhibit essential differences in their tuning as compared to direction cells; (ii) that anchor cells do not spuriously arise from potential collinearities between the factors “direction” and “anchor bearing” in our ANOVA framework; and (iii) that anchor cells can also be identified in an ANOVA framework with a reduced number of predictors (i.e., with factors “place” and “anchor bearing” instead of “direction”, “place”, and “anchor bearing”).

**Table S1.** Patients.

| **Patient** | **Sessions** | **Total number of trials** |
| --- | --- | --- |
| Freiburg_001 | Freiburg_001a | 39 |
|  | Freiburg_001b | 78 |
| Freiburg_002 | Freiburg_002 | 34 |
| Freiburg_003 | Freiburg_003a | 167 |
|  | Freiburg_003b | 162 |
| Freiburg_004 | Freiburg_004a | 166 |
|  | Freiburg_004b | 162 |
| Freiburg_005 | Freiburg_005 | 54 |
| Freiburg_006 | Freiburg_006 | 98 |
| Freiburg_007 | Freiburg_007 | 36 |
| Freiburg_008 | Freiburg_008 | 67 |
| Freiburg_009 | Freiburg_009 | 162 |
| Freiburg_010 | Freiburg_010 | 102 |
| Freiburg_011 | Freiburg_011 | 54 |
| Freiburg_012 | Freiburg_012 | 102 |
| Freiburg_013 | Freiburg_013a | 167 |
|  | Freiburg_013b | 164 |
| Freiburg_014 | Freiburg_014 | 94 |

**Table S2.**
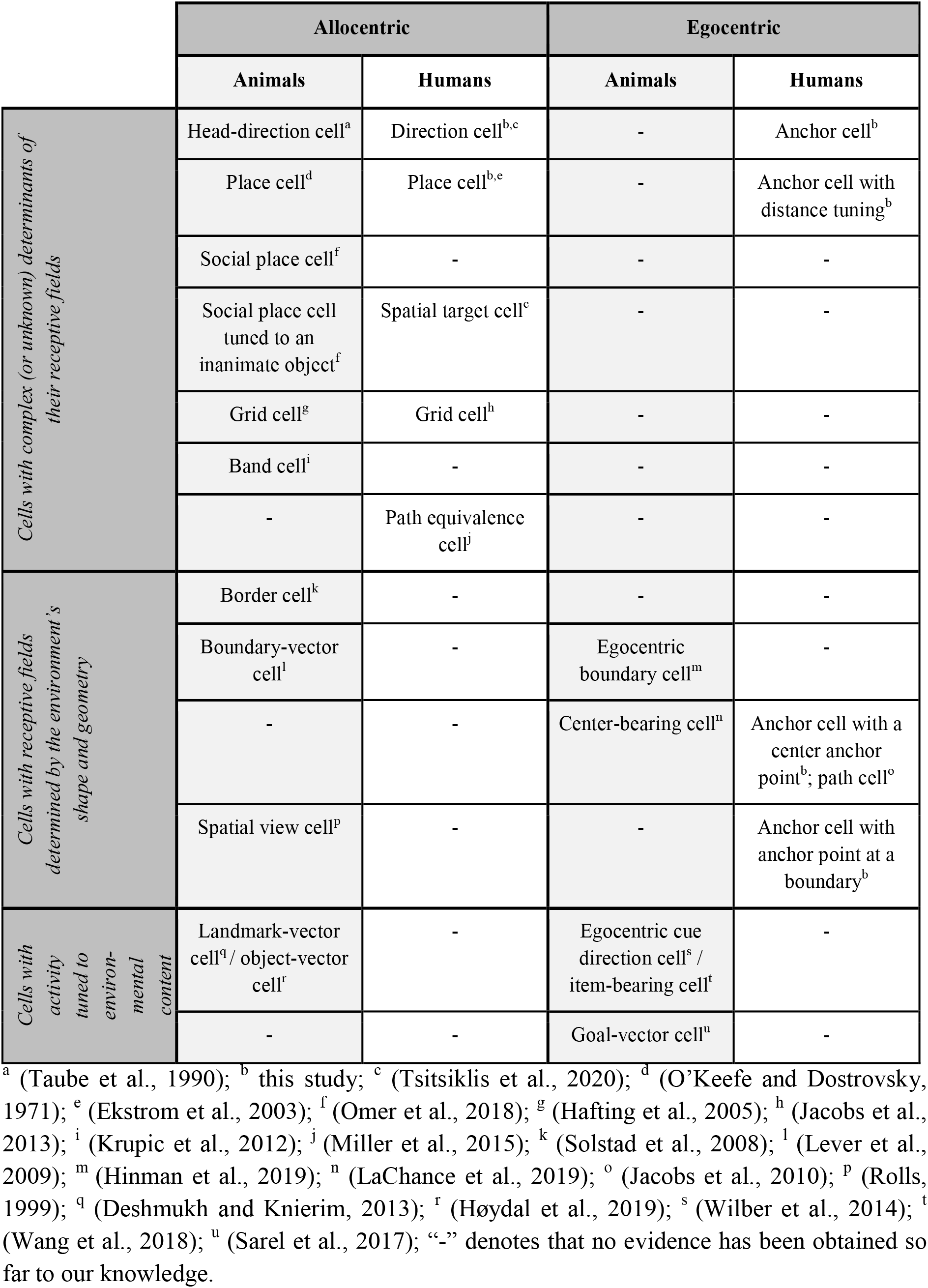
Allocentric and egocentric single-neuron codes in the medial temporal lobe.

|  | Allocentric |  | Egocentric |  |
| --- | --- | --- | --- | --- |
|  | Animals | Humans | Animals | Humans |
| *Cells with complex (or unknown) determinants of their receptive fields* | Head-direction cell<sup>a</sup> | Direction cell<sup>b,c</sup> | - | Anchor cell<sup>b</sup> |
|  | Place cell<sup>d</sup> | Place cell<sup>b,e</sup> | - | Anchor cell with distance tuning<sup>b</sup> |
|  | Social place cell<sup>f</sup> | - | - | - |
|  | Social place cell tuned to an inanimate object<sup>f</sup> | Spatial target cell<sup>c</sup> | - | - |
|  | Grid cell<sup>g</sup> | Grid cell<sup>h</sup> | - | - |
|  | Band cell<sup>i</sup> | - | - | - |
|  | - | Path equivalence cell<sup>j</sup> | - | - |
| *Cells with receptive fields determined by the environment's shape and geometry* | Border cell<sup>k</sup> | - | - | - |
|  | Boundary-vector cell<sup>l</sup> | - | Egocentric boundary cell<sup>m</sup> | - |
|  | - | - | Center-bearing cell<sup>n</sup> | Anchor cell with a center anchor point<sup>b</sup>; path cell<sup>o</sup> |
|  | Spatial view cell<sup>p</sup> | - | - | Anchor cell with anchor point at a boundary<sup>b</sup> |
| *Cells with activity tuned to environmental content* | Landmark-vector cell<sup>q</sup>/ object-vector cell<sup>r</sup> | - | Egocentric cue direction cell<sup>s</sup> / item-bearing cell<sup>t</sup> | - |
|  | - | - | Goal-vector cell<sup>u</sup> | - |

<sup>a</sup> (Taube et al., 1990); <sup>b</sup> this study; <sup>c</sup> (Tsitsiklis et al., 2020); <sup>d</sup> (O'Keefe and Dostrovsky, 1971); <sup>e</sup> (Ekstrom et al., 2003); <sup>f</sup> (Omer et al., 2018); <sup>g</sup> (Hafting et al., 2005); <sup>h</sup> (Jacobs et al., 2013); <sup>i</sup> (Krupic et al., 2012); <sup>j</sup> (Miller et al., 2015); <sup>k</sup> (Solstad et al., 2008); <sup>l</sup> (Lever et al., 2009); <sup>m</sup> (Hinman et al., 2019); <sup>n</sup> (LaChance et al., 2019); <sup>o</sup> (Jacobs et al., 2010); <sup>p</sup> (Rolls, 1999); <sup>q</sup> (Deshmukh and Knierim, 2013); <sup>r</sup> (Høydal et al., 2019); <sup>s</sup> (Wilber et al., 2014); <sup>t</sup> (Wang et al., 2018); <sup>u</sup> (Sarel et al., 2017); “-” denotes that no evidence has been obtained so far to our knowledge.

**Figure S1.**
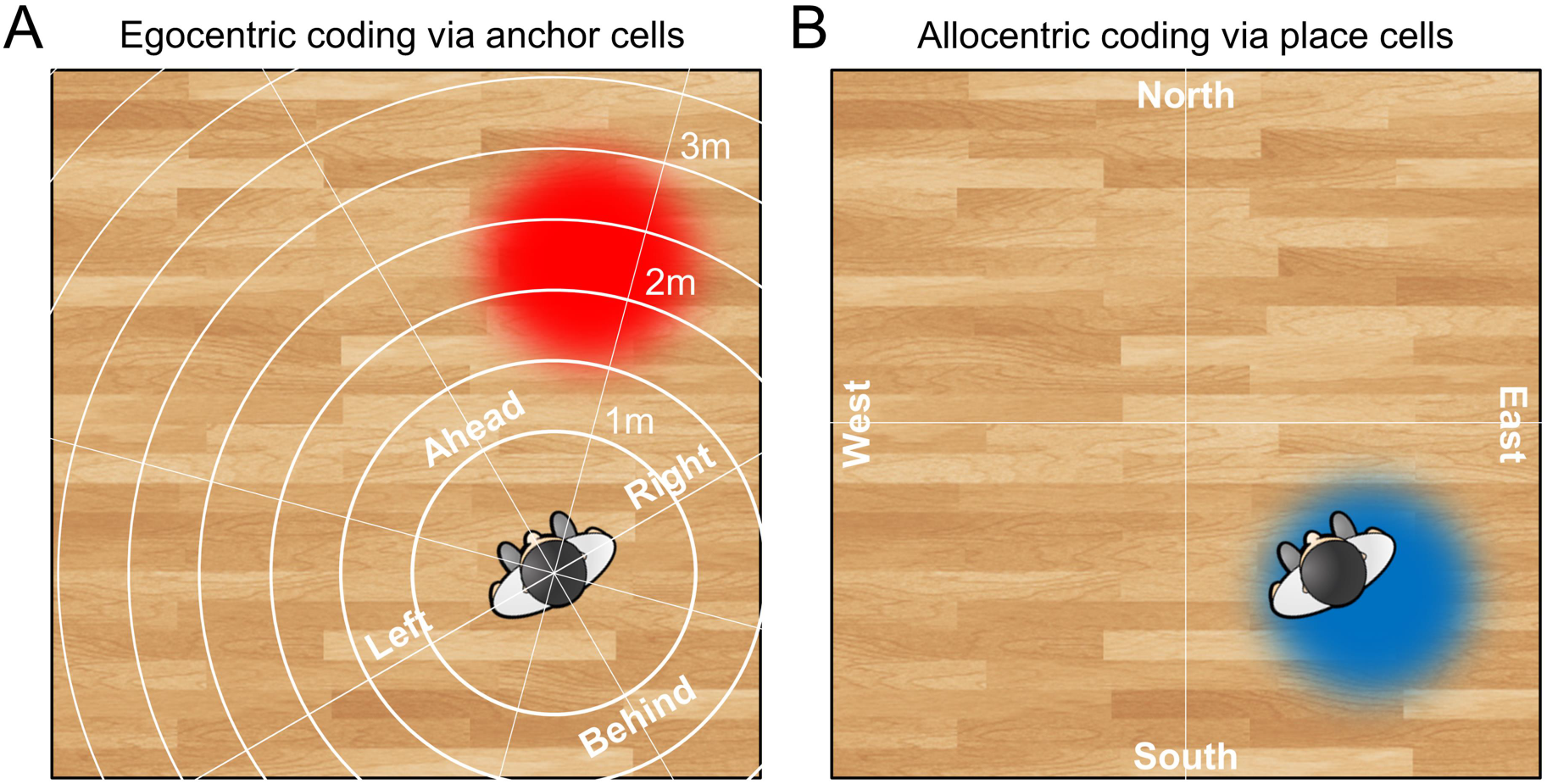
Illustration of the egocentric coding scheme of anchor cells as compared to the allocentric coding scheme of place cells. (**A**) Coding of spatial information in an egocentric reference frame (white lines), which is centered on the subject. The subject and the surrounding room are shown from a bird’s eye view. The anchor point of a hypothetical anchor cell is shown in red. The activity of this anchor cell provides the subject with the information that the area of the environment that is marked by the anchor point is about 45° to the right and about two meters away from the subject. (**B**) Coding of spatial information in an allocentric reference frame, which is bound to the external environment. The place field of a hypothetical place cell is shown in blue. The activity of this place cell provides the subject with the information that the subject is standing in the south-east part of the environment.

**Figure S2.**
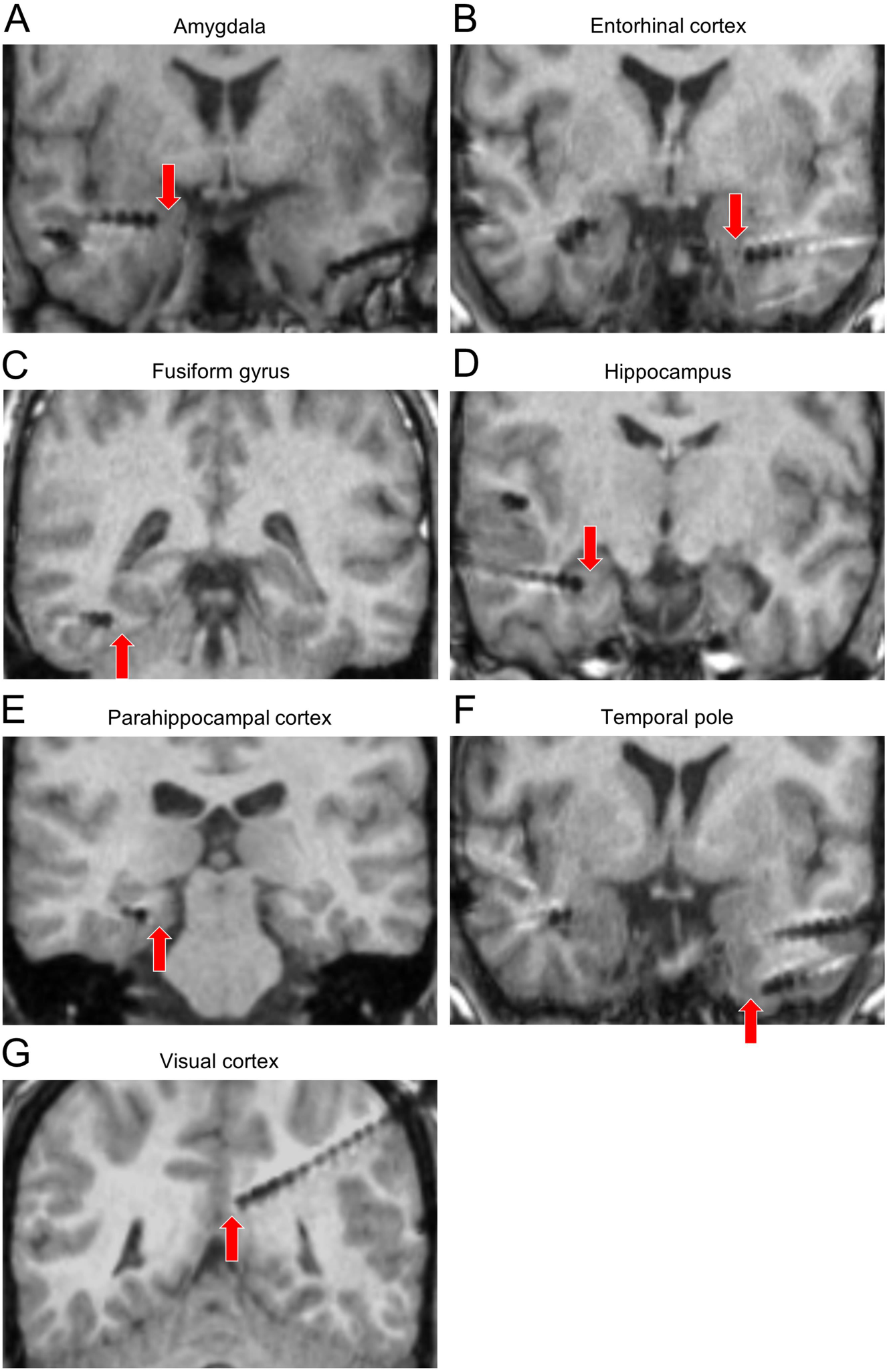
Examples of microelectrode locations. (**A to G**) Example microelectrode locations in amygdala, entorhinal cortex, fusiform gyrus, hippocampus, parahippocampal cortex, temporal pole, and visual cortex. Electrode contacts of depth electrodes appear as dark circles on the MRI scans. Red arrows point at putative microelectrode locations, which protrude 3-5 mm from the tip of the depth electrode (often not visible on MRI scans).

**Figure S3.**
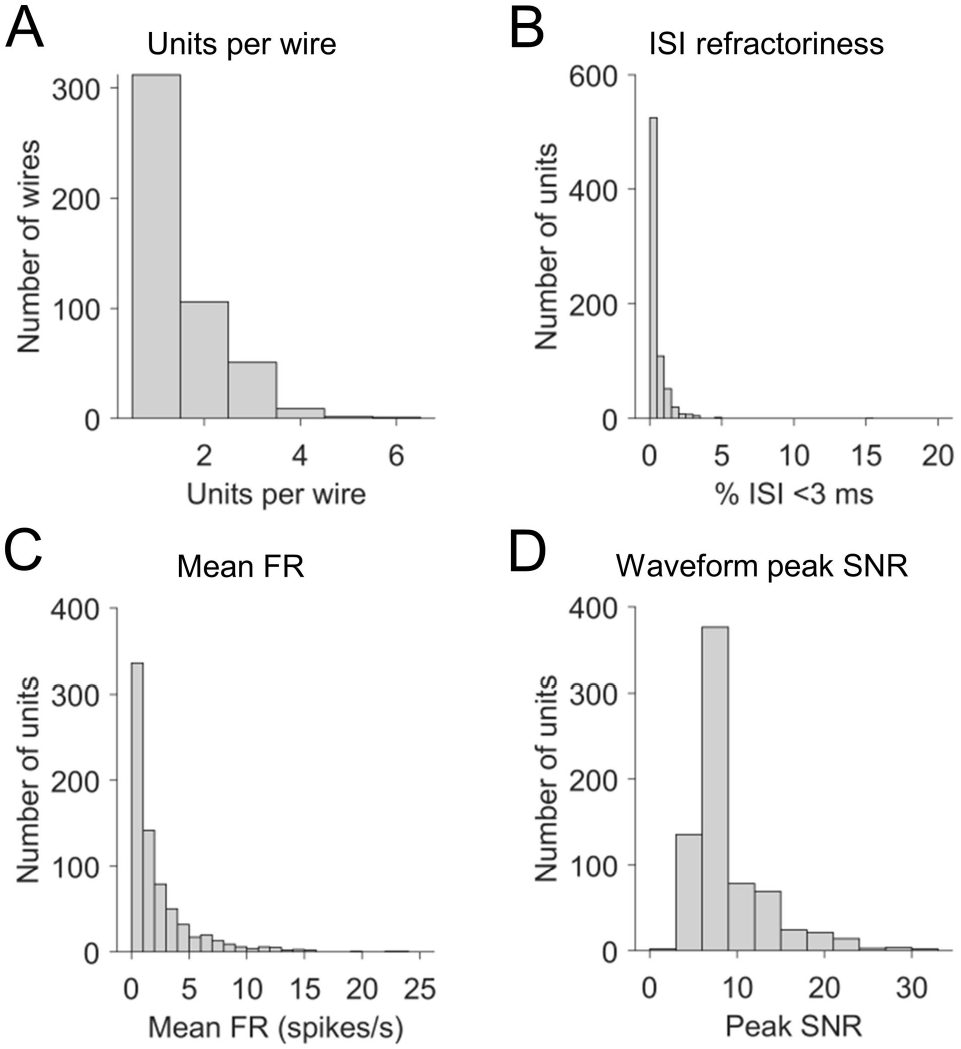
Quality assessment of neuronal recordings. (**A**) Histogram of units per wire. On average, 1.516 ± 0.037 units per wire (mean ± SEM) were recorded. (**B**) Histogram of the percentages of inter-spike intervals (ISIs) that were shorter than 3 ms. On average, units exhibited 0.434 ± 0.031% ISIs that were shorter than 3 ms (mean ± SEM). All units except for one had a value of <5%. (**C**) Histogram of mean firing rates (FRs). On average, units exhibited mean FRs of 2.268 ± 0.112 spikes/s (mean ± SEM). (**D**) Histogram of the mean waveform peak signal-to-noise ratio (SNR) of each unit. On average, the SNR of the mean waveform peak was 8.820 ± 0.168 (mean ± SEM), similar to previous reports [e.g., (Faraut et al., 2018)].

**Figure S4.**
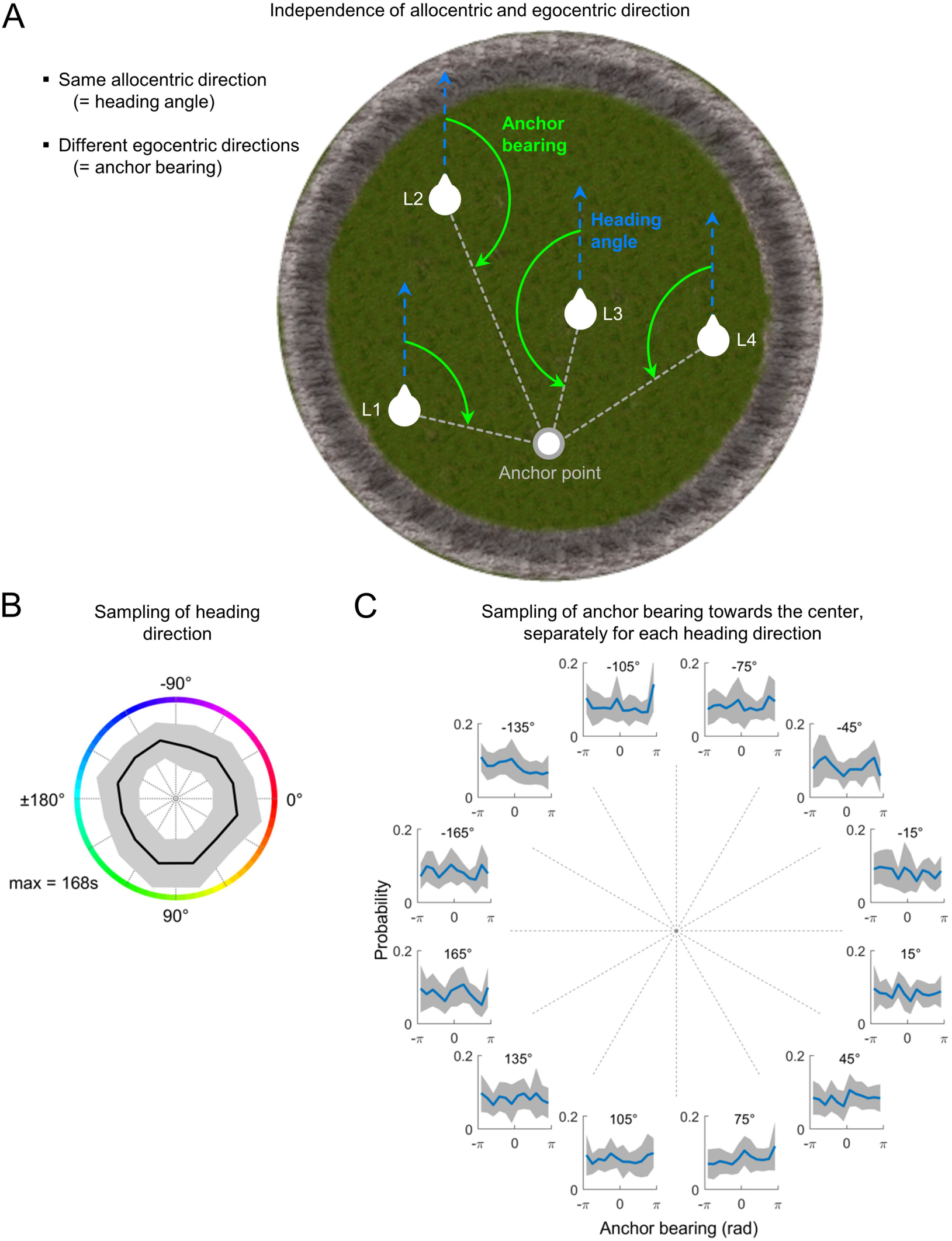
Sampling of egocentric and allocentric direction. (**A**) In allocentric coordinates, directional information is determined relative to the external world (for example, a specific location can be “south”). In egocentric coordinates, directional information is determined in relation to the self (for example, a specific location can be to the right of the subject). This diagram illustrates the general independence of egocentric anchor bearing and allocentric heading direction. A potential anchor point is depicted in the south part of the environment (gray circle). At different locations within the virtual environment (L1 to L4), the allocentric heading angle of the navigating subject is identical (light blue, dashed arrows), but the egocentric anchor bearings are different (light green arrows). White dashed lines, vectors between the subject’s locations and the anchor point. (**B**) Sampling of allocentric heading direction. Black line, mean across patients; gray area, SD across patients; max, maximum of direction-specific mean dwell times. (**C**) For each allocentric heading direction, the distribution of egocentric anchor bearings towards the environmental center is depicted, which is chosen as an example anchor point. Distributions are expressed as probabilities. A bearing of 0 corresponds to “anchor point ahead” and ±π corresponds to “anchor point behind”. Negative bearing values correspond to “anchor point to the left”. Blue line, mean across patients; gray area, SD across patients. Numbers above each subpanel indicate the allocentric heading direction for which the distribution of bearings is shown (with negative allocentric heading directions pointing north).

**Figure S5.**
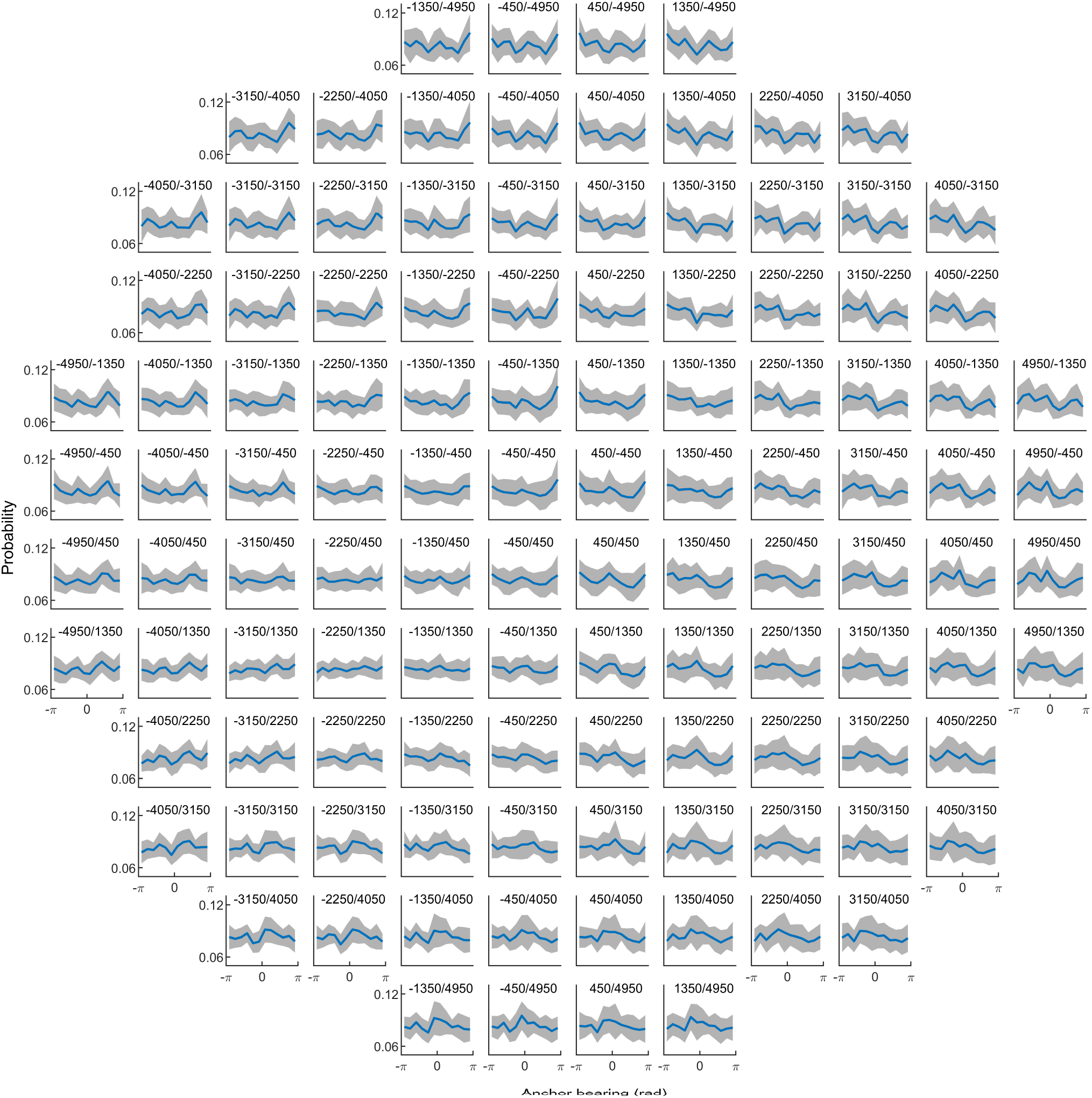
Sampling of egocentric bearing towards candidate anchor points. For each candidate anchor point, the distribution of bearings towards this candidate point is depicted. Distributions are expressed as probabilities. An anchor bearing of 0 corresponds to “anchor point ahead” and ±π corresponds to “anchor point behind”. Negative values correspond to “anchor point to the left”. Blue line, mean across patients. Gray area, SD across subjects. Numbers above each subpanel indicate the (x/y)-coordinate of the candidate anchor point (candidate points with negative y-values are located in the “north” part of the environment).

**Figure S6.**
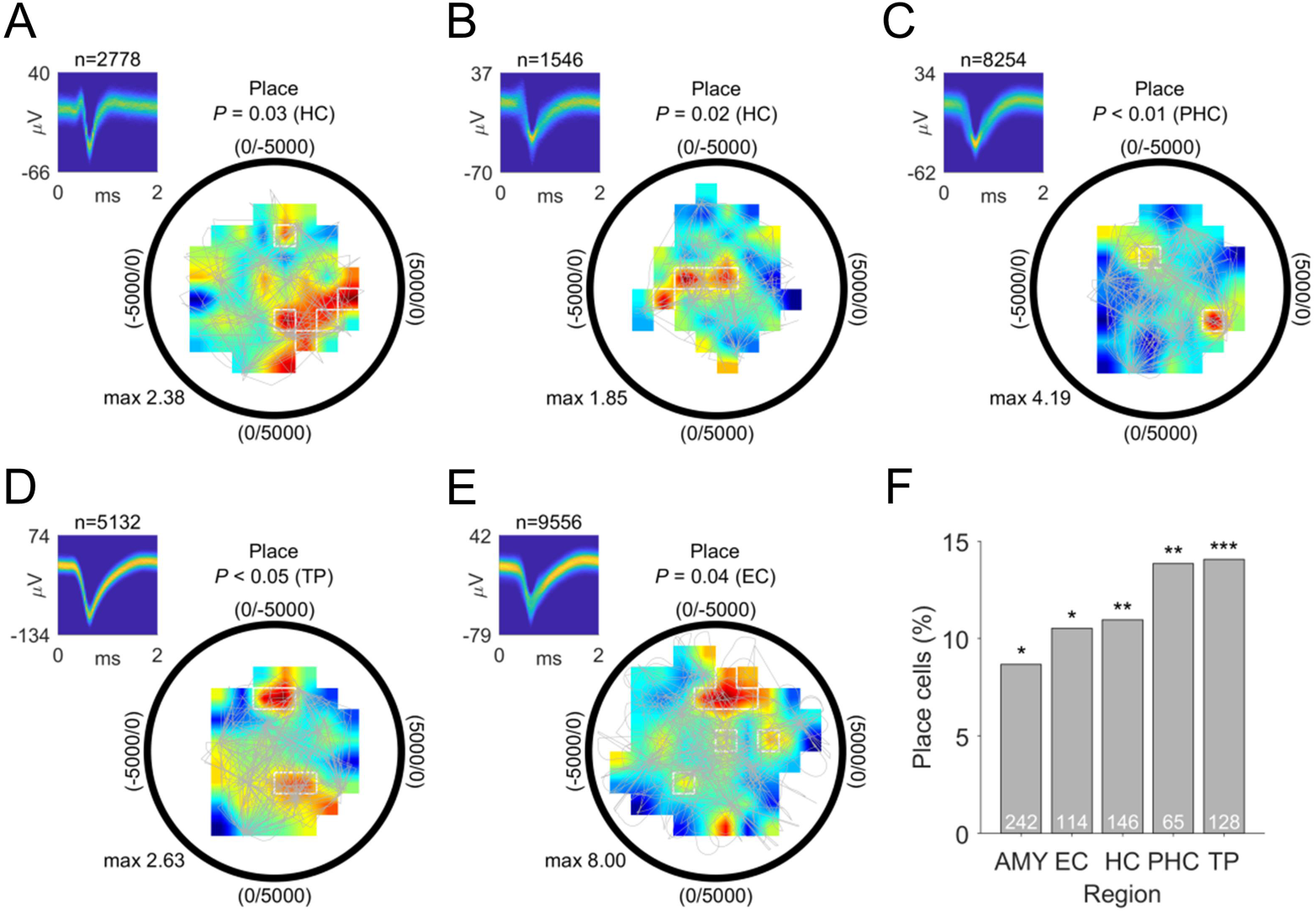
Place cells. (**A to E**) Examples of place cells. The hippocampal place cell shown in (**A**) increased its firing rate whenever the subject was in the southeast part of the environment. Colored areas depict smoothed firing rates as a function of place (dark blue, low firing rate; dark red, high firing rate); max, maximum firing rate (Hz). White line delineates place bins. Gray line, patient’s navigation path. Only bins with ≥5 separate traversals were included in the analysis to ensure sufficient behavioral sampling. Black circle, environmental boundary. Upper left subpanels show spike-density plots (number above subpanel indicates spike number entering the analysis); ms, milliseconds. (**F**) Distribution of place cells across brain regions (*n* = 84; binomial test, *P* < 0.001). AMY, amygdala; EC, entorhinal cortex; HC, hippocampus; PHC, parahippocampal cortex; TP, temporal pole. \**P* < 0.05; \*\**P* < 0.01; \*\*\**P* < 0.001.

## References

Acharya, L., Aghajan, Z.M., Vuong, C., Moore, J.J., and Mehta, M.R. (2016). Causal Influence of Visual Cues on Hippocampal Directional Selectivity. Cell 164, 197–207.

Aguirre, G.K., and D’Esposito, M. (1999). Topographical disorientation: a synthesis and taxonomy. Brain 122, 1613–1628.

Alexander, A.S., Carstensen, L.C., Hinman, J.R., Raudies, F., William Chapman, G., and Hasselmo, M.E. (2020). Egocentric boundary vector tuning of the retrosplenial cortex. Sci. Adv. 6, eaaz2322.

Aminoff, E.M., Kveraga, K., and Bar, M. (2013). The role of the parahippocampal cortex in cognition. Trends Cogn. Sci. 17, 379–390.

Barron, H.C., Vogels, T.P., Behrens, T.E., and Ramaswami, M. (2017). Inhibitory engrams in perception and memory. Proc. Natl. Acad. Sci. U. S. A. 114, 6666–6674.

Bastin, J., Vidal, J.R., Bouvier, S., Perrone-Bertolotti, M., Bénis, D., Kahane, P., David, O., Lachaux, J.-P., and Epstein, R.A. (2013). Temporal components in the parahippocampal place area revealed by human intracerebral recordings. J. Neurosci. 33, 10123–10131.

Bellmund, J.L.S., Gärdenfors, P., Moser, E.I., and Doeller, C.F. (2018). Navigating cognition: Spatial codes for human thinking. Science 362, eaat6766.

Berens, P. (2009). CircStat: A MATLAB Toolbox for Circular Statistics. J. Stat. Softw. 31, 1–21.

Bicanski, A., and Burgess, N. (2018). A neural-level model of spatial memory and imagery. Elife 7.

Binder, J.R., and Desai, R.H. (2011). The neurobiology of semantic memory. Trends Cogn. Sci. 15, 527–536.

Burgess, N. (2006). Spatial memory: how egocentric and allocentric combine. Trends Cogn. Sci. 10, 551–557.

Burgess, N. (2008). Spatial Cognition and the Brain. Ann. N. Y. Acad. Sci. 1124, 77–97.

Burgess, N., Cacucci, F., Lever, C., and O’Keefe, J. (2005). Characterizing multiple independent behavioral correlates of cell firing in freely moving animals. Hippocampus 15, 149–153.

Buzsáki, G., and Moser, E.I. (2013). Memory, navigation and theta rhythm in the hippocampal-entorhinal system. Nat. Neurosci. 16, 130–138.

Chaure, F.J., Rey, H.G., and Quian Quiroga, R. (2018). A novel and fully automatic spike-sorting implementation with variable number of features. J. Neurophysiol. 120, 1859–1871.

Chen, D., Kunz, L., Wang, W.X., Zhang, H., Wang, W.X., Schulze-Bonhage, A., Reinacher, P.C., Zhou, W., Liang, S., Axmacher, N., et al. (2018a). Hexadirectional Modulation of Theta Power in Human Entorhinal Cortex during Spatial Navigation. Curr. Biol. 28, 3310–3315.e4.

Chen, G., King, J.A., Lu, Y., Cacucci, F., and Burgess, N. (2018b). Spatial cell firing during virtual navigation of open arenas by head-restrained mice. Elife 7.

Coughlan, G., Laczó, J., Hort, J., Minihane, A.-M., and Hornberger, M. (2018). Spatial navigation deficits — overlooked cognitive marker for preclinical Alzheimer disease? Nat. Rev. Neurol. 14, 496–506.

Deshmukh, S.S., and Knierim, J.J. (2013). Influence of local objects on hippocampal representations: Landmark vectors and memory. Hippocampus 23, 253–267.

Doeller, C.F., King, J.A., and Burgess, N. (2008). Parallel striatal and hippocampal systems for landmarks and boundaries in spatial memory. Proc. Natl. Acad. Sci. U. S. A. 105, 5915–5920.

Doeller, C.F., Barry, C., and Burgess, N. (2010). Evidence for grid cells in a human memory network. Nature 463, 657–661.

Duvelle, É., Grieves, R.M., Hok, V., Poucet, B., Arleo, A., Jeffery, K.J., and Save, E. (2019). Insensitivity of place cells to the value of spatial goals in a two-choice flexible navigation task. J. Neurosci. 39, 2522–2541.

Ekstrom, A.D. (2015). Why vision is important to how we navigate. Hippocampus 25, 731–735.

Ekstrom, A.D., and Isham, E.A. (2017). Human spatial navigation: representations across dimensions and scales. Curr. Opin. Behav. Sci. 17, 84–89.

Ekstrom, A., Spiers, H., Bohbot, V., and Rosenbaum, R. (2018). Human spatial navigation (Princeton University Press).

Ekstrom, A.D., Kahana, M.J., Caplan, J.B., Fields, T.A., Isham, E.A., Newman, E.L., and Fried, I. (2003). Cellular networks underlying human spatial navigation. Nature 425, 184–188.

Ekstrom, A.D., Arnold, A.E.G.F., and Iaria, G. (2014). A critical review of the allocentric spatial representation and its neural underpinnings: toward a network-based perspective. Front. Hum. Neurosci. 8, 803.

Epstein, R., and Kanwisher, N. (1998). A cortical representation of the local visual environment. Nature 392, 598–601.

Epstein, R.A., Patai, E.Z., Julian, J.B., and Spiers, H.J. (2017). The cognitive map in humans: Spatial navigation and beyond. Nat. Neurosci. 20, 1504–1513.

Faraut, M.C.M., Carlson, A.A., Sullivan, S., Tudusciuc, O., Ross, I., Reed, C.M., Chung, J.M., Mamelak, A.N., and Rutishauser, U. (2018). Dataset of human medial temporal lobe single neuron activity during declarative memory encoding and recognition. Sci. Data 5, 180010.

Fried, I., Wilson, C.L., Maidment, N.T., Engel, J., Behnke, E., Fields, T.A., Macdonald, K.A., Morrow, J.W., and Ackerson, L. (1999). Cerebral microdialysis combined with single-neuron and electroencephalographic recording in neurosurgical patients. J. Neurosurg. 91, 697–705.

Gallistel, C.R. (1990). The Organization of Learning.

Gauthier, J.L., and Tank, D.W. (2018). A Dedicated Population for Reward Coding in the Hippocampus. Neuron 99, 179–193.e7.

Gofman, X., Tocker, G., Weiss, S., Boccara, C.N., Lu, L., Moser, M.-B., Moser, E.I., Morris, G., and Derdikman, D. (2019). Dissociation between Postrhinal Cortex and Downstream Parahippocampal Regions in the Representation of Egocentric Boundaries. Curr. Biol. 29, 2751–2757.e4.

Hafting, T., Fyhn, M., Molden, S., Moser, M.-B., and Moser, E.I. (2005). Microstructure of a spatial map in the entorhinal cortex. Nature 436, 801–806.

Hardcastle, K., Maheswaranathan, N., Ganguli, S., and Giocomo, L.M. (2017). A Multiplexed, Heterogeneous, and Adaptive Code for Navigation in Medial Entorhinal Cortex. Neuron 94.

Hartigan, J.A., and Hartigan, P.M. (1985). The Dip Test of Unimodality. Ann. Stat. 13, 70–84.

Hinman, J.R., Chapman, G.W., and Hasselmo, M.E. (2019). Neuronal representation of environmental boundaries in egocentric coordinates. Nat. Commun. 10, 2772.

Høydal, Ø.A., Skytøen, E.R., Andersson, S.O., Moser, M.-B., and Moser, E.I. (2019). Object-vector coding in the medial entorhinal cortex. Nature 568, 400–404.

Ismakov, R., Barak, O., Jeffery, K., and Derdikman, D. (2017). Grid Cells Encode Local Positional Information. Curr. Biol. 27, 2337–2343.e3.

Jacobs, J., Kahana, M.J., Ekstrom, A.D., Mollison, M. V, and Fried, I. (2010). A sense of direction in human entorhinal cortex. Proc. Natl. Acad. Sci. U. S. A. 107, 6487–6492.

Jacobs, J., Weidemann, C.T., Miller, J.F., Solway, A., Burke, J.F., Wei, X.-X., Suthana, N., Sperling, M.R., Sharan, A.D., Fried, I., et al. (2013). Direct recordings of grid-like neuronal activity in human spatial navigation. Nat. Neurosci. 16, 1188–1190.

Jercog, P.E., Ahmadian, Y., Woodruff, C., Deb-Sen, R., Abbott, L.F., and Kandel, E.R. (2019). Heading direction with respect to a reference point modulates place-cell activity. Nat. Commun. 10, 2333.

Klatzky, R.L. (1998). Allocentric and Egocentric Spatial Representations: Definitions, Distinctions, and Interconnections. (Springer, Berlin, Heidelberg), pp. 1–17.

Krupic, J., Burgess, N., and O’Keefe, J. (2012). Neural representations of location composed of spatially periodic bands. Science (80-.). 337, 853–857.

Kunz, L., Schröder, T.N., Lee, H., Montag, C., Lachmann, B., Sariyska, R., Reuter, M., Stirnberg, R., Stöcker, T., Messing-Floeter, P.C., et al. (2015). Reduced grid-cell-like representations in adults at genetic risk for Alzheimer’s disease. Science 350, 430–433.

Kunz, L., Maidenbaum, S., Chen, D., Wang, L., Jacobs, J., and Axmacher, N. (2019a). Mesoscopic Neural Representations in Spatial Navigation. Trends Cogn. Sci. 23, 615–630.

Kunz, L., Wang, L., Lachner-Piza, D., Zhang, H., Brandt, A., Dümpelmann, M., Reinacher, P.C., Coenen, V.A., Chen, D., Wang, W.-X., et al. (2019b). Hippocampal theta phases organize the reactivation of large-scale electrophysiological representations during goal-directed navigation. Sci. Adv. 5, eaav8192.

Kutter, E.F., Bostroem, J., Elger, C.E., Mormann, F., and Nieder, A. (2018). Single Neurons in the Human Brain Encode Numbers. Neuron 100, 753–761.e4.

LaChance, P.A., Todd, T.P., and Taube, J.S. (2019). A sense of space in postrhinal cortex. Science 365, eaax4192.

Lever, C., Burton, S., Jeewajee, A., O’Keefe, J., and Burgess, N. (2009). Boundary vector cells in the subiculum of the hippocampal formation. J. Neurosci. 29, 9771–9777.

Manns, J.R., Howard, M.W., and Eichenbaum, H. (2007). Gradual Changes in Hippocampal Activity Support Remembering the Order of Events. Neuron 56, 530–540.

Meilinger, T., and Vosgerau, G. (2010). Putting Egocentric and Allocentric into Perspective. (Springer, Berlin, Heidelberg), pp. 207–221.

Miller, J., Watrous, A.J., Tsitsiklis, M., Lee, S.A., Sheth, S.A., Schevon, C.A., Smith, E.H., Sperling, M.R., Sharan, A., Asadi-Pooya, A.A., et al. (2018). Lateralized hippocampal oscillations underlie distinct aspects of human spatial memory and navigation. Nat. Commun. 9, 2423.

Miller, J.F., Neufang, M., Solway, A., Brandt, A., Trippel, M., Mader, I., Hefft, S., Merkow, M., Polyn, S.M., Jacobs, J., et al. (2013). Neural activity in human hippocampal formation reveals the spatial context of retrieved memories. Science 342, 1111–1114.

Miller, J.F., Fried, I., Suthana, N., and Jacobs, J. (2015). Repeating Spatial Activations in Human Entorhinal Cortex. Curr. Biol. 25, 1080–1085.

Mormann, F., Kornblith, S., Cerf, M., Ison, M.J., Kraskov, A., Tran, M., Knieling, S., Quian Quiroga, R., Koch, C., and Fried, I. (2017). Scene-selective coding by single neurons in the human parahippocampal cortex. Proc. Natl. Acad. Sci. U. S. A. 114, 1153–1158.

Moser, E.I., Moser, M.B., and McNaughton, B.L. (2017). Spatial representation in the hippocampal formation: A history. Nat. Neurosci. 20.

O’Keefe, J., and Dostrovsky, J. (1971). The hippocampus as a spatial map. Preliminary evidence from unit activity in the freely-moving rat. Brain Res. 34, 171–175.

Olson, J.M., Li, J.K., Montgomery, S.E., and Nitz, D.A. (2020). Secondary Motor Cortex Transforms Spatial Information into Planned Action during Navigation. Curr. Biol. 0.

Omer, D.B., Maimon, S.R., Las, L., and Ulanovsky, N. (2018). Social place-cells in the bat hippocampus. Science 359, 218–224.

Oostenveld, R., Fries, P., Maris, E., and Schoffelen, J.-M. (2011). FieldTrip: Open source software for advanced analysis of MEG, EEG, and invasive electrophysiological data. Comput. Intell. Neurosci. 2011, 156869.

Pfeiffer, B.E., and Foster, D.J. (2013). Hippocampal place-cell sequences depict future paths to remembered goals. Nature 497, 74–79.

Ploner, C.J., Gaymard, B.M., Rivaud-Péchoux, S., Baulac, M., Clémenceau, S., Samson, S., and Pierrot-Deseilligny, C. (2000). Lesions Affecting the Parahippocampal Cortex Yield Spatial Memory Deficits in Humans. Cereb. Cortex 10, 1211–1216.

Qasim, S.E., Miller, J., Inman, C.S., Gross, R.E., Willie, J.T., Lega, B., Lin, J.-J., Sharan, A., Wu, C., Sperling, M.R., et al. (2019). Memory retrieval modulates spatial tuning of single neurons in the human entorhinal cortex. Nat. Neurosci. 433862.

Reber, T.P., Bausch, M., Mackay, S., Boström, J., Elger, C.E., and Mormann, F. (2019). Representation of abstract semantic knowledge in populations of human single neurons in the medial temporal lobe. PLOS Biol. 17, e3000290.

Rolls, E.T. (1999). Spatial view cells and the representation of place in the primate hippocampus. Hippocampus 9, 467–480.

Sarel, A., Finkelstein, A., Las, L., and Ulanovsky, N. (2017). Vectorial representation of spatial goals in the hippocampus of bats. Science 355, 176–180.

Schacter, D.L., Addis, D.R., and Buckner, R.L. (2007). Remembering the past to imagine the future: The prospective brain. Nat. Rev. Neurosci. 8, 657–661.

Shahi, M., Dhingra, S., Sandler, R., Rios, R., Vuong, C., Acharya, L., and Mehta, M.R. (2019). Hippocampal anchor fields. Soc. Neurosci. Annu. Meet.

Shuman, T., Aharoni, D., Cai, D.J., Lee, C.R., Chavlis, S., Page-Harley, L., Vetere, L.M., Feng, Y., Yang, C.Y., Mollinedo-Gajate, I., et al. (2020). Breakdown of spatial coding and interneuron synchronization in epileptic mice. Nat. Neurosci. 23, 229–238.

Solstad, T., Boccara, C.N., Kropff, E., Moser, M.-B., and Moser, E.I. (2008). Representation of geometric borders in the entorhinal cortex. Science 322, 1865–1868.

Staresina, B.P., and Wimber, M. (2019). A Neural Chronometry of Memory Recall. Trends Cogn. Sci. 23, 1071–1085.

Taube, J.S., Muller, R.U., and Ranck, J.B. (1990). Head-direction cells recorded from the postsubiculum in freely moving rats. I. Description and quantitative analysis. J. Neurosci. 10, 420–435.

Tsitsiklis, M., Miller, J., Qasim, S.E., Inman, C.S., Gross, R.E., Willie, J.T., Smith, E.H., Sheth, S.A., Schevon, C.A., Sperling, M.R., et al. (2020). Single-Neuron Representations of Spatial Targets in Humans. Curr. Biol. 30, 245–253.e4.

Waller, D., and Hodgson, E. (2006). Transient and enduring spatial representations under disorientation and self-rotation. J. Exp. Psychol. Learn. Mem. Cogn. 32, 867–882.

Wang, R.F., and Spelke, E.S. (2002). Human spatial representation: Insights from animals. Trends Cogn. Sci. 6, 376–382.

Wang, C., Chen, X., Lee, H., Deshmukh, S.S., Yoganarasimha, D., Savelli, F., and Knierim, J.J. (2018). Egocentric coding of external items in the lateral entorhinal cortex. Science 362, 945–949.

Wang, C., Chen, X., and Knierim, J.J. (2020). Egocentric and allocentric representations of space in the rodent brain. Curr. Opin. Neurobiol. 60, 12–20.

Weniger, G., and Irle, E. (2006). Posterior parahippocampal gyrus lesions in the human impair egocentric learning in a virtual environment. Eur. J. Neurosci. 24, 2406–2414.

Wilber, A.A., Clark, B.J., Forster, T.C., Tatsuno, M., and McNaughton, B.L. (2014). Interaction of egocentric and world-centered reference frames in the rat posterior parietal cortex. J. Neurosci. 34, 5431–5446.

Wood, E.R., Dudchenko, P.A., Robitsek, R.J., and Eichenbaum, H. (2000). Hippocampal Neurons Encode Information about Different Types of Memory Episodes Occurring in the Same Location. Neuron 27, 623–633.

Zhang, H., Zherdeva, K., and Ekstrom, A.D. (2014). Different “routes” to a cognitive map: dissociable forms of spatial knowledge derived from route and cartographic map learning. Mem. Cogn. 42, 1106–1117.

